# The pigtail macaque (*Macaca nemestrina*) model of COVID-19 reproduces diverse clinical outcomes and reveals new and complex signatures of disease

**DOI:** 10.1101/2021.08.28.458047

**Authors:** Alexandra Melton, Lara A Doyle-Meyers, Robert V Blair, Cecily Midkiff, Hunter J Melton, Kasi Russell-Lodrigue, Pyone P Aye, Faith Schiro, Marissa Fahlberg, Dawn Szeltner, Skye Spencer, Brandon J Beddingfield, Kelly Goff, Nadia Golden, Toni Penney, Breanna Picou, Krystle Hensley, Kristin E Chandler, Jessica A Plante, Kenneth S Plante, Scott C Weaver, Chad J Roy, James A Hoxie, Hongmei Gao, David C Montefiori, Joseph L Mankowski, Rudolf P Bohm, Jay Rappaport, Nicholas J Maness

## Abstract

The novel coronavirus SARS-CoV-2, the causative agent of COVID-19 disease, has killed over four million people worldwide as of July 2021 with infections rising again due to the emergence of highly transmissible variants. Animal models that faithfully recapitulate human disease are critical for assessing SARS-CoV-2 viral and immune dynamics, for understanding mechanisms of disease, and for testing vaccines and therapeutics. Pigtail macaques (PTM, *Macaca nemestrina*) demonstrate a rapid and severe disease course when infected with simian immunodeficiency virus (SIV), including the development of severe cardiovascular symptoms that are pertinent to COVID-19 manifestations in humans. We thus proposed this species may likewise exhibit severe COVID-19 disease upon infection with SARS-CoV-2. Here, we extensively studied a cohort of SARS-CoV-2-infected PTM euthanized either 6- or 21-days after respiratory viral challenge. We show that PTM demonstrate largely mild-to-moderate COVID-19 disease. Pulmonary infiltrates were dominated by T cells, including CD4+ T cells that upregulate CD8 and express cytotoxic molecules, as well as virus-targeting T cells that were predominantly CD4+. We also noted increases in inflammatory and coagulation markers in blood, pulmonary pathologic lesions, and the development of neutralizing antibodies. Together, our data demonstrate that SARS-CoV-2 infection of PTM recapitulates important features of COVID-19 and reveals new immune and viral dynamics and thus may serve as a useful animal model for studying pathogenesis and testing vaccines and therapeutics.

## Introduction

In late 2019, a novel coronavirus was found circulating in humans in China. This virus showed substantial genomic similarities with the severe acute respiratory syndrome coronavirus (SARS-CoV) that caused an outbreak and panic in 2003^1^ in addition to a number of bat sarbecoviruses^2^; hence, it was named SARS-CoV-2^3^. SARS-CoV-2 is the causative agent of COVID-19 disease and a worldwide pandemic that has killed more than four million persons to date including over 600,000 deaths in the United States.

Though most infected individuals exhibit no or mild symptoms, a subset experience severe complications, including highly elevated pro-inflammatory cytokines and coagulation biomarkers, acute respiratory distress syndrome (ARDS), and death^4–9^. Most available data suggest that the intensity of the immune response plays a role in determining COVID-19 severity and progression, with severe disease occurring approximately 3-to-4-weeks after initial symptoms.^10, 11^. Thus, a deep understanding of the immunopathologic mechanisms of disease in those with advanced disease and of viral clearance in asymptomatic infection and those with mild disease is critical for the development of next generation therapies and vaccines.

Animal models that faithfully recapitulate human disease are needed to assess the roles of particular cell subsets in disease etiology. Various species of nonhuman primates can be infected by SARS-CoV-2 and exhibit disease ranging from mild to severe^12–16^. The use of timed infections with well characterized viral stocks in animals with relatively high genetic similarity with humans allows the dissection of immune responses with nuance and detail not possible in humans. The most widely used species of NHP for COVID-19 research has been the rhesus macaque (*Macaca mulatta*). This model has proved valuable for testing vaccines as viral infection dynamics in this species are robust and well-studied and therefore can be compared between treatment groups. However, SARS-CoV-2-induced disease in this species is generally mild and does not recapitulate the more severe disease seen in a subset of humans. Thus, multiple NHP models are needed to capture the spectrum of disease seen in humans. In this study, we infected pigtail macaques (PTM, *Macaca nemestrina*) with SARS-CoV-2 (WA1/2020 isolate) to assess this novel animal model of COVID-19 disease.

PTM are a unique and valuable animal model for other viral diseases. Simian immunodeficiency virus (SIV) infection of rhesus macaques (RhM) is the most widely used nonhuman primate model of HIV/AIDS and is used widely for testing vaccines and cure strategies. However, SIV-associated disease in RhM can take up to several years to develop, somewhat limiting their use for studying disease mechanisms. In contrast, infection of PTM with the same viral isolates leads to rapid disease development with enhanced cardiovascular manifestations relative to RhM, which is of particular relevance to COVID-19 disease^17–20^. Thus, we proposed that SARS-CoV-2 infection of PTM may likewise lead to accelerated COVID-19 disease or demonstrate immune features of disease not detected in other animal models. If so, this species will be valuable for assessing COVID-19 disease mechanisms and for testing novel vaccines and therapeutics. We tracked viral and immune dynamics through the course of infection in a cohort of PTM. We found that disease in this model largely mirrored that observed in RhM but with unique immune features, such as pulmonary infiltration of CD4+ T cells that exhibit antiviral and cytotoxic functions, as is seen in COVID-19 patients^21^.

Together, our data characterize, in depth, a novel animal model that may prove useful for assessing moderate COVID-19 disease mechanisms and testing new therapeutics.

## Materials and Methods

### Animal cohort, viral inoculations, and procedures

Four male pigtail macaques (PTM), between the ages of 5 and 6 years old (Table S1), were exposed to 1×10^6^ TCID_50_ of SARS-CoV-2 USA WA1/2020 (World Reference Center for Emerging Viruses and Arboviruses, Galveston, TX) through both intranasal and intratracheal inoculation. The viral stock was sequenced and determined to have no mutations at greater than 5% of reads that differed from the original patient isolate. Pre- and post-exposure samples included blood, bronchoalveolar lavage (BAL), and mucosal swabs (nasal, pharyngeal, rectal, and bronchial brush). Physical examination and imaging (radiography Figure S1) were conducted before viral exposure and weekly after exposure. Animals were monitored for 6 (n=2) or 21 (n=2) days before euthanasia and tissue harvest. At necropsy, samples from each of the major lung lobes (left and right, cranial, middle, and caudal lobes) were collected in TRIzol (Invitrogen, Lithuania) and fresh frozen at −80°C. The remainder of the lung lobes were infused and then immersed in formalin fixative. The rest of the necropsy was performed routinely with collection of tissues from all major organs in DMEM media, fresh frozen, or in formalin fixative.

| Animal ID | Sex | Age (y) | Weight (kg) |
| --- | --- | --- | --- |
| MA27 | Male | 6.07 | 7.55 |
| MA30 | Male | 5.61 | 5.80 |
| MA24 | Male | 5.64 | 8.40 |
| MA28 | Male | 5.81 | 8.60 |

### Ethics Statement

Pigtail macaques used in this study were purpose bred at the University of Washington National Primate Research Center for experiments. Macaques were housed in compliance with the NRC Guide for the Care and Use of Laboratory Animals and the Animal Welfare Act. Animal experiments were approved by the Institutional Animal Care and Use Committee of Tulane University (protocol P0451). The Tulane National Primate Research Center (TNPRC) is fully accredited by AAALAC International, Animal Welfare Assurance No. A3180-01. During the study, animals were singly housed indoors in climate-controlled conditions with a 12/12-light/dark cycle. All the animals on this study were monitored twice daily to ensure their welfare. Any abnormalities, including those of appetite, stool, and behavior, were recorded and reported to a veterinarian. The animals were fed commercially prepared nonhuman primate diet twice daily. Supplemental foods were provided in the form of fruit, vegetables, and foraging items as part of the TNPRC environmental enrichment program. Water was available ad libitum through an automatic watering system. The TNPRC environmental enrichment program is reviewed and approved by the IACUC semi-annually. Veterinarians in the TNPRC Division of Veterinary Medicine have established procedures to minimize pain and distress using several approaches. Animals were anesthetized with ketamine-HCl (10 mg/kg) or tiletamine/zolazepam (3-8 mg/kg) prior to all procedures. Preemptive and post procedural analgesia (buprenorphine 0.01 mg/kg or buprenorphine sustained-release 0.2 mg/kg SQ) was used for procedures that would likely cause more than momentary pain or distress in humans undergoing the same procedures. The above listed anesthetics and analgesics were used to minimize pain and distress in accordance with the recommendations of the Weatherall Report. The animals were euthanized at the end of the study using methods consistent with recommendations of the American Veterinary Medical Association (AVMA) Panel on euthanasia and per the recommendations of the IACUC. Specifically, the animals were anesthetized with tiletamine/zolazepam (8 mg/kg IM) and given buprenorphine (0.01 mg/kg IM) followed by an overdose of pentobarbital sodium. Death was confirmed using auscultation to confirm the cessation of respiratory and circulatory functions and by the lack of corneal reflexes.

### Isolation of Viral RNA

The *Quick*-RNA Viral Kit (Zymo Research, Irvine, CA) was used to isolate viral RNA (vRNA) from mucosal swab and bronchial brush samples collected in 200 µL DNA/RNA Shield 1X (Zymo Research, Irvine, CA) following the manufacturer’s protocol. Briefly, 400 µL DNA/RNA viral buffer was added to the swab samples. In a modification to the manufacturer’s protocol, swabs were transferred directly to the Zymo spin column for centrifugation. The vRNA was eluted in 50 µL elution buffer.

### Viral RNA Quantification by Quantitative Real-Time PCR

Quantification of viral RNA was performed as described^22^ using the CDC N1 primers/probe for quantification of total viral RNA and with primers/probe specific for the nucleocapsid subgenomic RNA to provide an estimate of replicating virus. Specifically, vRNA was quantified using the QuantStudio 6 Real-Time PCR System (Applied Biosystems, Waltham, MA). Five microliters vRNA was added in duplicate to a 0.1 mL 96-well MicroAmp fast optical reaction plate (Applied Biosystems, REF# 4346906). For genomic vRNA quantification, the 2019-nCoV RUO Kit (Integrated DNA Technologies, Coralville, IA) was used, according to the manufacturer’s protocol, to target the N1 amplicon of the N gene along with TaqPath 1-Step RT-qPCR Master Mix (Applied Biosystems Waltham, MA). For the subgenomic assay, a forward primer targeting the subgenomic leader sequence and a reverse primer/probe (Integrated DNA Technologies, Waltham, MA) designed to target the N gene, was used along with the TaqPath Master Mix mentioned above. Fifteen microliters of the respective master mix was added to each well and run using the following conditions: 25°C for 2 minutes, 50°C for 15 minutes, 95°C for 2 minutes followed by 40 cycles of 95°C for 3 seconds and 60°C for 30 seconds. In vitro transcribed RNA was quantified and diluted to known copy numbers and used to generate the genomic and subgenomic standard curves. Both genomic and subgenomic viral copy numbers were calculated by plotting Cq values from unknown samples against the respective standard curve. Positive, negative, and non-template controls were analyzed along with each set of samples.

### Isolation of PBMCs

Peripheral blood mononuclear cells (PBMCs) were isolated from whole blood using SepMate-50 Isolation tubes (Stem Cell Technologies, Vancouver, Canada) per the manufacturer’s protocol. Cells were counted using a Cellometer Auto 2000 (Nexcelom, Lawrence, MA), resuspended in Bambanker cell freezing medium (GC Lymphotec, Tokyo, Japan) at approximately 1×10^7^ cells/mL and cryopreserved at −80°C.

### ELISA assays

D-dimer levels in sodium citrate plasma samples were measured via an enzyme-linked immunosorbent assay (ELISA) (Ray Biotech, Peachtree Corners, GA) per the manufacturer’s protocol. Samples were diluted 600,000-fold and plated in duplicate.

Plates were analyzed using the GloMax Explorer plate reader (Promega, Madison, WI) and GraphPad Prism (GraphPad Software version 9, LaJolla, California). Heatmap was generated using Microsoft Excel. Data was normalized by dividing raw data values from Day 6, 14 and 21 by the baseline value for each animal.

Kynurenine and tryptophan levels in plasma were measured using commercially available enzyme-linked immunosorbent assays (Rocky Mountain Diagnostics, Colorado Springs, CO) per the manufacturer’s protocol. The GloMax Explorer plate reader (Promega, Madison, WI) along with GraphPad Prism v9 were used to analyze the plates.

### Quantification of Inflammatory Cytokines and Coagulation Biomarkers

BioLegend’s bead-based immunosorbent assays were used to measure inflammatory cytokines in serum (LegendPlex NHP Inflammation Panel, BioLegend, San Diego, CA) and coagulation biomarkers in sodium citrate plasma (LegendPlex Human Fibrinolysis Panel). Serum and plasma samples were diluted 4-fold and 10,000-fold, respectively, and assayed in duplicate. Results were read using a MacsQuant 16 Flow Cytometer (Miltenyi Biotec) and LegendPlex’s online data analysis tool (Qognit). Heatmap was generated using Microsoft Excel. Data was normalized by dividing raw data values from Day 6, 14 and 21 by the baseline value for each animal.

### Flow Cytometry Analysis

Phenotypic and intracellular cytokine analysis of mononuclear cells (MNC) isolated from blood and bronchoalveolar lavage (BAL) was performed using antibodies against markers listed in Supplementary Tables 2-4. Briefly, cells were washed and counted with the Cellometer Auto 2000 (Nexcelom Bioscience, Lawrence, MA). Cells were then pelleted and resuspended in Live/Dead stain cocktail (50 µL PBS + 0.5 µL live/dead stain per test) (Fixable Aqua Dead Cell Stain Kit, Invitrogen, Lithuania) and incubated in the dark for 20 minutes. Cells were washed in PBS supplemented with 2% FBS, pelleted, resuspended, and incubated in surface-stain cocktail consisting of 50µL BD Horizon Brilliant Violet Stain Buffer (BD Bioscience, Franklin Lakes, NJ) plus antibodies (see Supplementary Tables 2-4) for 20 minutes in the dark. Cells were washed in PBS with 2% FBS, pelleted, then resuspended in 200 µL BD Cytofix/Cytoperm solution (BD Biosciences, Franklin Lakes, NJ) and incubated in the dark for 20 minutes. Cells were washed in BD Perm/Wash Buffer (BD Biosciences, Franklin Lakes, NJ), pelleted, and resuspended in intracellular-staining cocktail consisting of 100 µL BD Perm/Wash Buffer plus antibodies according to Supplementary Tables 2-4 and incubated for 20 minutes in the dark. Finally, cells were then washed, pelleted, and resuspended in 200 µL 1x BD Stabilizing Fixative (BD Biosciences, Franklin Lakes, NJ).

### Monocyte Cytokine Expression

To measure monocyte cytokine expression, MNCs from blood and BAL were washed and counted (Cellometer Auto 2000, Nexcelom Bioscience, Lawrence, MA), pelleted, and then resuspended in DMEM (Gibco, Grand Island, NY) with 5% Anti-Anti (Gibco, Grand Island, NY) at 1×10^6^ cells/mL. Cells were stimulated with lipopolysaccharide at 10 ng/mL (Sigma, St Louis, MO) and incubated with 1 µL/mL Brefeldin-A (BioLegend, San Diego, CA) for 4-6 hours at 37°C, 5% CO_2_. Cells were then stained following the procedure described above with antibodies listed in the Monocyte Panel (Table S2).

### T cell Cytokine Expression

MNCs from BAL were counted, washed, pelleted, and resuspended in DMEM with 5% Anti-Anti at 1×10^6^ cells/mL. T cell cytokine expression was measured by stimulating MNCs with cell stimulation cocktail (Biolegend, San Diego, CA) for 4-6 hours at 37°C, 5% CO_2_. To measure T cell responses to SARS-CoV-2 antigens, MNCs from blood and BAL were washed, pelleted and resuspended in DMEM with 5% Anti-Anti and 10% FBS at 1×10^6^ cells/mL followed by overnight stimulation at 37°C, 5% CO_2_ with either cell stimulation cocktail or with one of the following viral peptide pools obtained through BEI Resources, NIAID, NIH: Peptide Array, SARS Coronavirus Nucleocapsid Protein (NR-52419), Spike Glycoprotein (NR-52402), Membrane Protein (NR-53822), or Envelope Protein (NR-53822) along with Brefeldin-A. Cells were stained as described above using the antibodies listed in the T cell panel (Table S4).

All samples were acquired on a LSRFortessa Cell Analyzer (BD Biosciences, Franklin Lakes, NJ) using BD FACSDIVA 8.0.1 software. Approximately 1×10^6^ cells were acquired from each sample. Data was analyzed using FlowJo version 10.7.1 for MAC (Becton Dickinson and Company, Ashland, OR). SARS-CoV-2 antigen specific T cell response figures (9A&B) were generated using the Matlab based tool cyt3^23^. Data was transformed using arcsin 150. Cytokine expression was measured in FlowJo and, when applicable, applied to cyt3 generated figures. t-distributed stochastic neighbor embedding (tSNE) analysis was performed in FlowJo 10.7.1, nightingale rose plots were generated in R using the ggplot2 package, radial plots were generated in Microsoft Excel.

### Histopathology and Immunohistochemistry

Zinc-formalin fixed tissues were processed routinely, embedded in paraffin and cut in to 5 µm sections for hematoxylin and eosin (H&E) or immunohistochemical (IHC) staining. For H&E staining, tissue samples were collected in Zinc formalin (Anatech, Sparks, NV) and immersion fixed for a minimum of 72 hours before being washed and dehydrated using a Thermo Excelsior AS processor. Upon removal from the processor, tissues were transferred to a Thermo Shandon Histocentre 3 embedding station where they were submersed in warm paraffin and allowed to cool into blocks. From these blocks, 5um sections were cut and mounted on charged glass slides, baked overnight at 60°C and passed through Xylene, graded ethanol, and double distilled water to remove paraffin and rehydrate tissue sections. A Leica Autostainer XL was used to complete the deparaffinization, rehydration and routine hematoxylin and eosin stain preparing the slides for examination by a board-certified veterinary pathologist using HALO software (Indica Labs, Albuquerque, NM).

For IHC staining, tissue sections were mounted on Superfrost Plus Microscope slides (Fisher Scientific, Carlsbad, CA), incubated for 1 hour at 60°C, and passed through Xylene, graded ethanol, and double distilled water to remove paraffin and rehydrate tissue sections. A microwave was used for heat induced epitope retrieval (HIER). Slides were boiled for 20 minutes in a Tris based solution, pH 9 (Vector Laboratories, Burlingame, CA), supplemented with 0.01% Tween-20. Slides were briefly rinsed in hot, distilled water and transferred to a hot citrate based solution, pH 6.0 (Vector Laboratories, Burlingame, CA) where they were allowed to cool to room temperature. All slide manipulation from this point forward was done at room temperature with incubations taking place in a black humidifying chamber. Once cool, slides were rinsed in tris buffered saline (TBS) and incubated with Background Punisher (Biocare Medical, Pacheco, CA) for 10 minutes. Slides were then submerged in a solution of TBS supplemented with 0.01% Triton×100 (TBS-T×100) and placed on a rocker platform for two 5 minute washes followed by a TBS rinse before being returned to humidifying chamber to be incubated with serum free protein block (Dako, Santa Clara, CA) for 20 minutes. Mouse anti-Granzyme primary antibody (Table S6) was then added to the slides and allowed to bind for 60 minutes. Slides were then washed twice with TBS-T×100 and once with TBS. The labeling of the antibody for visualization was done using the MACH3 AP kit (Biocare Medical, Pacheco, CA). Both the MACH3 probe and polymer were incubated for 20 minutes with TBS-T×100 and TBS washes in between. Slides were incubated with permanent red substrate (Dako, Santa Clara, CA) for 20 minutes and placed in TBS to halt the enzymatic reaction.

All other staining was done consecutively with the following method. Slides were incubated with a blocking buffer comprised of 10% normal goat serum (NGS) and 0.02% fish skin gelatin in phosphate buffered saline (PBS) for 40 minutes. This blocking buffer was also used to dilute both primary and secondary antibodies (Table S6). Primary antibodies were added to slides for 60 minutes. After washing two times with PBS supplemented with 0.02% fish skin gelatin and 0.01% Triton×100 (PBS-FSG-T×100) and once with PBS-FSG, slides were incubated for 40 minutes with a secondary antibody made in goat, raised against the primary host species, and tagged with an Alexa Fluor fluorochrome (488 or 568). The 3 washes mentioned above were repeated before DAPI nuclear stain was added for 10 minutes. Slides were mounted using anti-quenching mounting media containing Mowiol (Sigma, St Louis, MO) and DABCO (Sigma, St Louis, MO) and allowed to dry overnight before imaging with a Axio Slide Scanner (Zeiss, Hamburg, Germany). HALO software (Indica Labs Albuquerque, NM) was used for quantification and analysis.

### Detection of Neutralizing Antibodies in Serum

Pseudovirus neutralization testing of serum samples was performed using a SARS-CoV-2.D614G spike-pseudotyped virus in 293/ACE2 cells, with neutralization assessed via reduction in luciferase activity as described^24, 25^.

### Statistical Analysis

GraphPad Prism (version 9 GraphPad Software, LaJolla California) was used for all graphing and statistical analyses. The Kruskal-Wallis with Dunn’s multiple comparison test was used to compare changes in cell frequencies and surface marker, cytokine and Granzyme B expression,

## Results

### Viral dynamics

Four pigtail macaques (PTM) inoculated with SARS-CoV-2 were followed via blood, mucosal swab and bronchoalveolar lavage (BAL) sampling. Two animals were euthanized at 6 days post infection (dpi) and two at 21 dpi (Figure 1A). Quantitative RT-PCR was used to track viral genomic and subgenomic RNA through the course of the study at several sites. We detected both genomic and subgenomic SARS-CoV-2 RNA in all four animals throughout the first several days of infection (Figure 1B-K). One animal, MA27, euthanized at 6-days post infection (dpi), showed a spike in genomic and subgenomic viral RNA (vRNA) at necropsy in the pharynx (Figure 1D,E), with viral levels also beginning to rise in the nasal cavity (Figure 1B,C). MA28, euthanized at 21-dpi, showed detectable levels of vRNA in the nasal and rectal mucosa throughout the course of the study (Figure 1B,J). These findings are consistent with viral kinetics seen in Rhesus macaques with the possible exception of lower viral loads in the pharynx in the PTM model.

**Figure 1.**
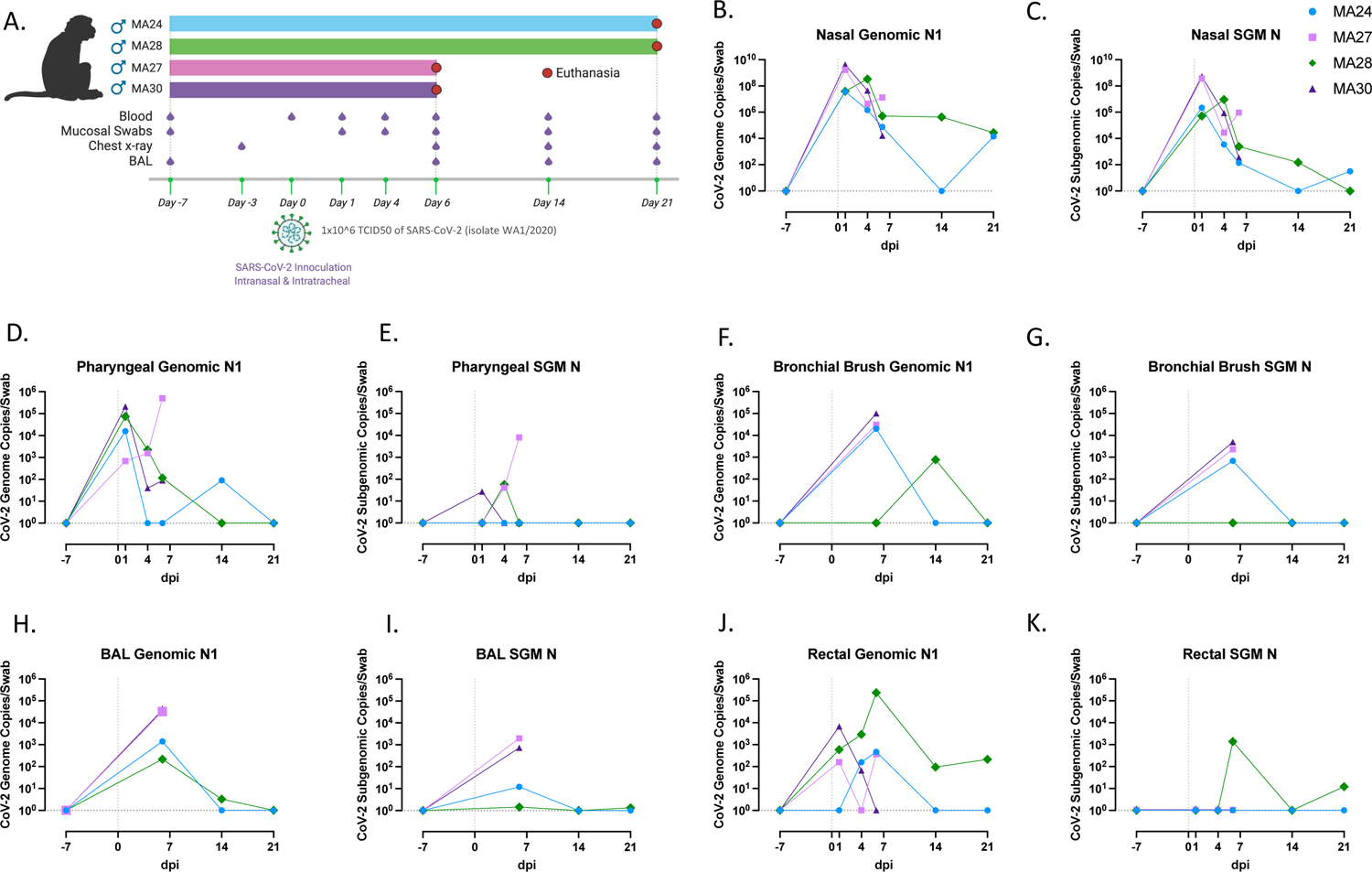
Viral dynamics. **A.** Outline of study design. Four PTM were exposed to 1×10^6 TCID50 of SARS-CoV-2 (isolate WA1/2020) through a combination of intranasal and intratracheal inoculation on Day 0. Figure created with BioRender.com. **B-K**. Quantification of SARS-CoV-2 RNA levels from mucosal swabs overtime (Quantitative RT PCR). Genomic (**B,D,F,H,J**) Subgenomic (**C,E,G,I,K**). B-K Baseline: n=4, Day 1: n=4, Day 4: n=4, Day 6: n=4, Day 14: n=2, Day 21: n=2

### Pulmonary disease and pathology

Thoracic radiographs were obtained from all animals before infection and weekly after, revealing subtle changes consistent with interstitial pneumonia reflective of mild to moderate COVID-19 (Figure S1A-L). Postmortem examination at 6-days post infection (dpi) revealed mild-to-moderate SARS-CoV-2-associated pneumonia in one of the two animals, MA27. The pneumonia was characterized by multifocal tan-plum areas of consolidation in the caudal left lung lobe (Figure S2A, Table S1). At 21-dpi, gross lesions were minimal and only observed in one of two animals, MA28. The lesions in this animal were two small, flat tan foci on the dorsolateral aspect of the left caudal lung lobe (Figure S2D, Table S1).

Histopathological findings consistent with SARS-CoV-2 associated pneumonia were observed in both animals at 6-dpi. Both animals had an interstitial pneumonia that was localized to regions of the left caudal lung. Regions of interstitial pneumonia were characterized by alveolar septa that were mild to markedly expanded by a mixture of macrophages, lymphocytes, and neutrophils. Alveolar septa were frequently lined by type II pneumocytes (Figure 2C&D), and alveoli contained large numbers of alveolar macrophages with rafts of fibrin in more severely affected areas (Figure 2A-D).

**Figure 2.**
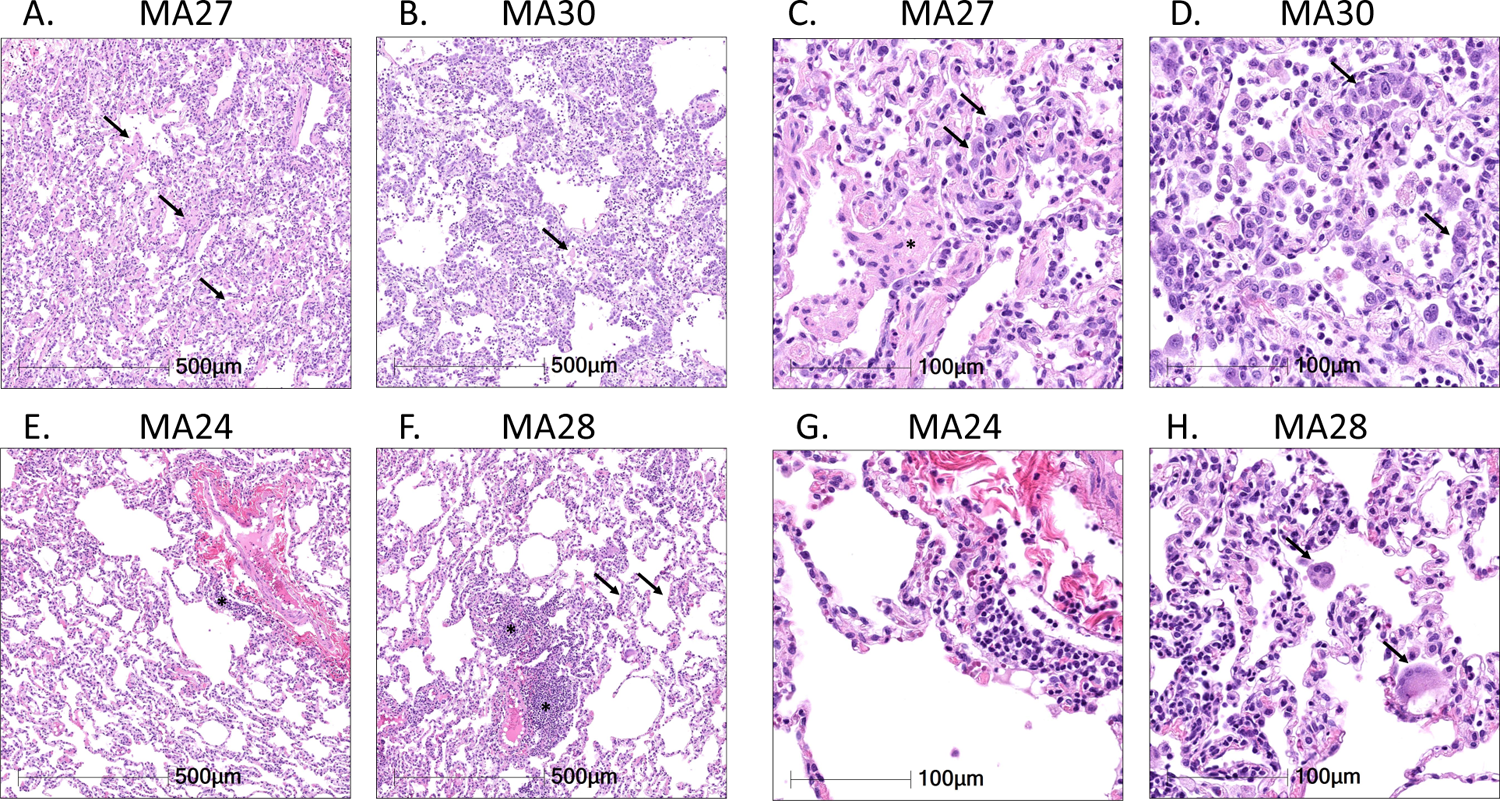
Histopathologic findings in SARS-CoV-2 infected pigtail macaques (PTM). Histopathologic findings at 6-**(A-D)** and 21-dpi **(E-H)**. **A&B.** At 6-dpi alveolar septa are expanded by inflammatory infiltrate and alveoli contain rafts of fibrin (**arrows**). **C&D.** The inflammatory infiltrate is composed of a mixture of histiocytes, lymphocytes, and neutrophils, and alveolar septa are frequently lined by type II pneumocytes (**arrows**). In severely affected areas, alveoli contain fibrin rafts (**C, asterisks**). **E&F.** At 21-dpi, there is residual inflammation composed of perivascular lymphoid aggregates (**asterisks**), and mild thickening of alveolar septa (**arrows). G&H**. The residual inflammation is composed predominately of lymphocytes, and in MA28, rare multinucleated giant cells (**H, arrows**).

At 21-dpi, minimal-to-mild residual interstitial pulmonary inflammation was observed in both animals. The residual inflammation was composed of perivascular lymphoid aggregates along with mild thickening of alveolar septa (Figure 2E&F). The inflammatory infiltrate at this time point was composed predominately of lymphocytes; however, in one animal, MA28, low numbers of multinucleated giant cells were present in alveoli. (Figure 2H).

### Serum cytokine measures of inflammation

We next measured a panel of cytokines in blood serum after infection. Fluctuations in several inflammatory cytokine levels, as compared to baseline, were found throughout the study. Interleukin-8 (IL-8), a neutrophil chemoattractant, was the most consistently increased cytokine at 6-dpi whereas IL-6 and IL-12-p40 decreased in all animals at day 6 (Figure 3A). Interestingly, MA27 had a stronger inflammatory cytokine response at 6-dpi compared to the other three animals, as exemplified by increases in several cytokines, including IL-10, IFN-γ, GM-CSF, IL-8, IL-17A, MCP-1 and most notably, TNF-⍺ and IFN-β. As stated previously, this animal had increasing viral loads at 6-dpi suggesting a possible link between the intensity of the inflammatory response and the level of replicating virus. Animal MA28, which exhibited consistently high genomic vRNA levels in both nasal and rectal swabs through 21-dpi, showed a rise IL-10, IL-1β, IL-12p40 and IP-10 serum levels at necropsy (21-dpi).

**Figure 3.**
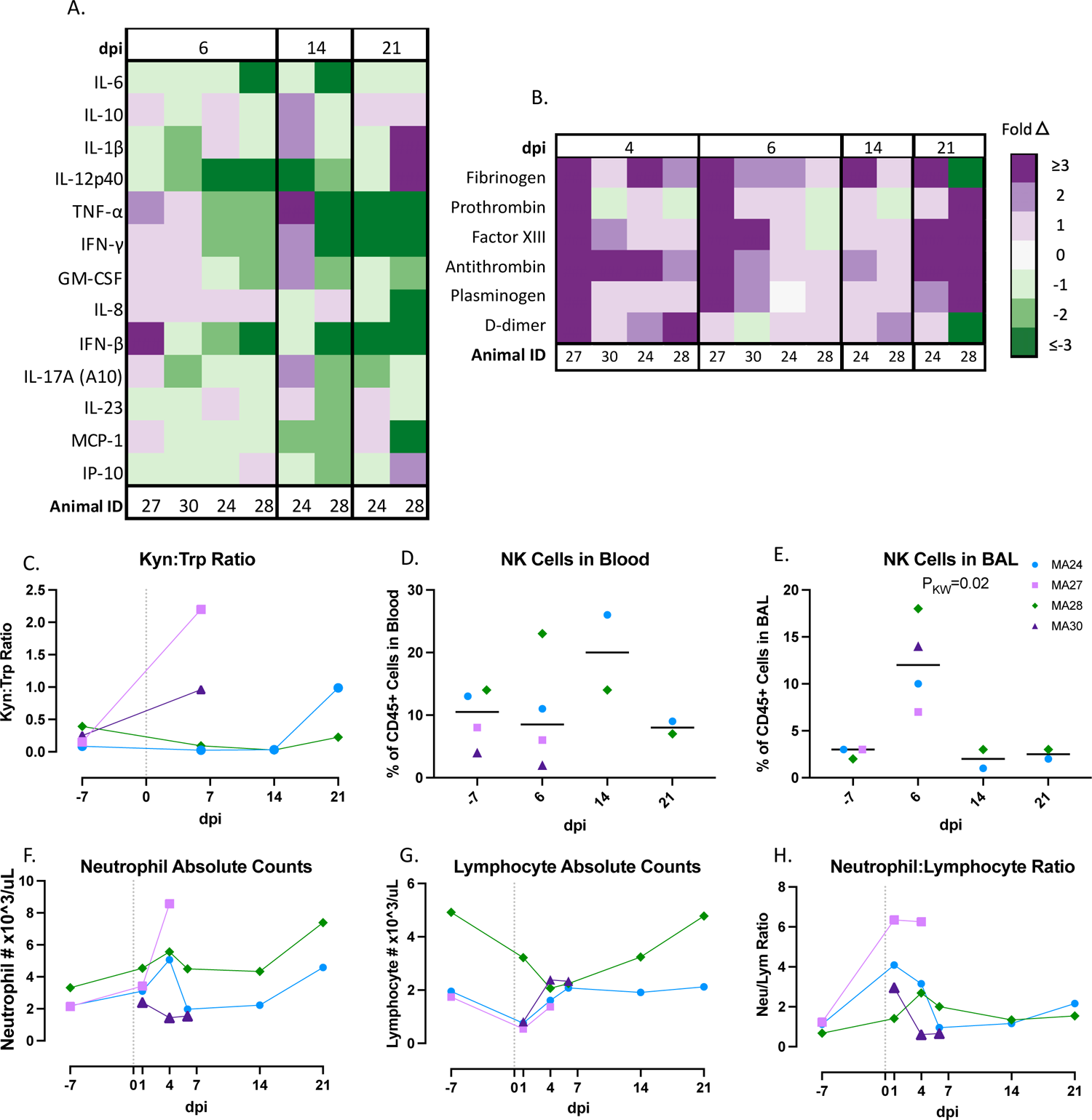
Inflammatory innate immune response in pigtail macaques challenged with SARS-CoV-2. **A.** Changes in serum cytokine levels at 6-, 14- and 21-days post SARS-CoV-2 infection. Data represent fold changes from baseline. **B.** Changes in coagulation biomarkers in plasma at 4-, 6-, 14- and 21-days post infection (dpi). Data are fold changes from baseline. **C.** Ratio of Kynurenine (Kyn) to Tryptophan (Trp) as a measure of indoleamine 2,3-dioxygenase (IDO) activity before and after SARS-CoV-2 infection. **D&E.** Frequency of Natural Killer (NK, CD45+ CD3-CD8+) cells in the blood **A.** (D) or **(E)** BAL at baseline, 6-, 14- and 21-days post infection (dpi). Bars represent median **F.** Absolute number of neutrophils pre- and post-SARS-CoV-2 infection. **G.** Absolute number of lymphocytes pre- and post-SARS-CoV-2 infection. **H.** Changes in neutrophil to lymphocyte ratio before and after SARS-CoV-2 infection. Figures 3C&D Baseline: n=4, Day 6: n=4, Day 14: n=2, Day 21: n=2; Figure 3E Baseline: n=3, Day 6: n=4, Day 14: n=2, Day 21: n=2; Figures 3F-H Baseline: n=3, Day 1: n=4, Day 4: n=4 Day 6: n=4, Day 14: n=2, Day 21: n=2. Figures C-H Day 0=day of infection. Kruskal-Wallis test for variance for overall medians used to determine significance. Kruskal-Wallis comparison of overall means (P_KW_) test used to determine significance. P values ≤0.05 reported

### Markers of coagulopathy

Complications related to coagulopathy have been reported in humans with severe COVID-19 disease, with highly elevated levels of D-dimers in particular a biomarker of disease severity^26, 27^. To examine whether PTM recapitulate this phenotype, we measured multiple biomarkers of coagulation in blood (Figure 3B), including fibrinogen, prothrombin, factor XIII, antithrombin, plasminogen, and D-dimers. We found nearly universal increases in coagulation biomarkers in the first week of infection. Specifically, we noted increased D-dimer levels in all four animals at 6-dpi, with MA27 and MA28 exhibiting a greater than 3-fold increase relative to baseline before resolving to near baseline levels. Interestingly, several biomarkers (fibrinogen, prothrombin, factor XIII, antithrombin, and plasminogen) began to rise again at 21-dpi.

### Kynurenine tryptophan pathway

Pro-inflammatory cytokines, specifically interferon gamma-γ (IFN-γ), promote the kynurenine (Kyn) pathway (KP) of tryptophan (Trp) catabolism^28^. Recent studies in humans hospitalized with COVID-19 suggest that the Kyn:Trp ratio positively correlates with disease severity^29^. We measured the Kyn:Trp ratio in plasma at baseline, and days 6, 14 and 21 (Figure 3C, S3A,B). Again, MA27 showed the greatest increase in the Kyn:Trp ratio at 6-dpi possibly providing another biomarker of the more severe disease course seen in this animal.

### NK cells

The initial immune response to SARS-CoV-2 infection involves the intricate interplay between cells of the innate immune system. Natural killer (NK) cells are cytotoxic lymphocytes that often play a key role in the early defense against viral infections.

Studies of hospitalized COVID-19 patients show that decreases in circulating NK cells correlate with disease severity^30, 31^. Here, we measured the percentage of NK cells (defined as CD45+ CD3-CD8+) in both the blood and bronchoalveolar lavage fluid (BAL) at baseline, and days 6-, 14-, and 21-post infection (Figure 3D&E). We did not find significant changes in NK cells in our study. However, MA28 and MA24 had slight increases in circulating NK cells at day 6 and day 14, respectively. Flow cytometry analysis of BAL indicated an increase in infiltrating NK cells in the lung at 6-dpi in all four animals.

### Neutrophil to lymphocyte ratio

A high incidence of neutrophilia coupled with lymphocytopenia has been reported in COVID-19 patients^31, 32^. Animals MA24, MA27, and MA28 all experienced neutrophilia and lymphocytopenia during the course of the study but the changes were mild and values largely stayed within normal limits. Pre-infection data on these cells were not available for MA30 (Table S1). The neutrophil to lymphocyte ratio (NLR) has been identified as an important predictor of disease severity in human patients^33^. We thus measured the NLR at baseline and 6-, 14- and 21-dpi. MA27 had the highest NLR at 1- and 4-days post infection (Figure 3F-H). These data are consistent with the increased inflammatory cytokine and D-dimer levels, higher K:T ratio, and increasing viral titers at the time of necropsy observed in MA27. Each potentially correlate with or contribute to the more severe lung pathology observed in this animal at necropsy.

### SARS-CoV-2 infection and macrophage pulmonary infiltration

Fluorescent immunohistochemistry of the lung for SARS-CoV-2 nucleoprotein identified small clusters of SARS-CoV-2 infected cells, predominately lining the alveolar septa, in both animals sacrificed at 6-dpi (Figure 4 A,B&E). COVID-19 disease is commonly characterized by pulmonary infiltration of inflammatory immune cells^34^. Innate cells, particularly monocytes/macrophages are considered important mediators of disease progression^35^. At 6-dpi, the alveoli contained large numbers of IBA1+ macrophages (Figures 4A-D,F). By 21-dpi, macrophage numbers were greatly reduced and no SARS-CoV-2+ cells were detected in either MA24 or MA28 (Figure 4C-F).

**Figure 4.**
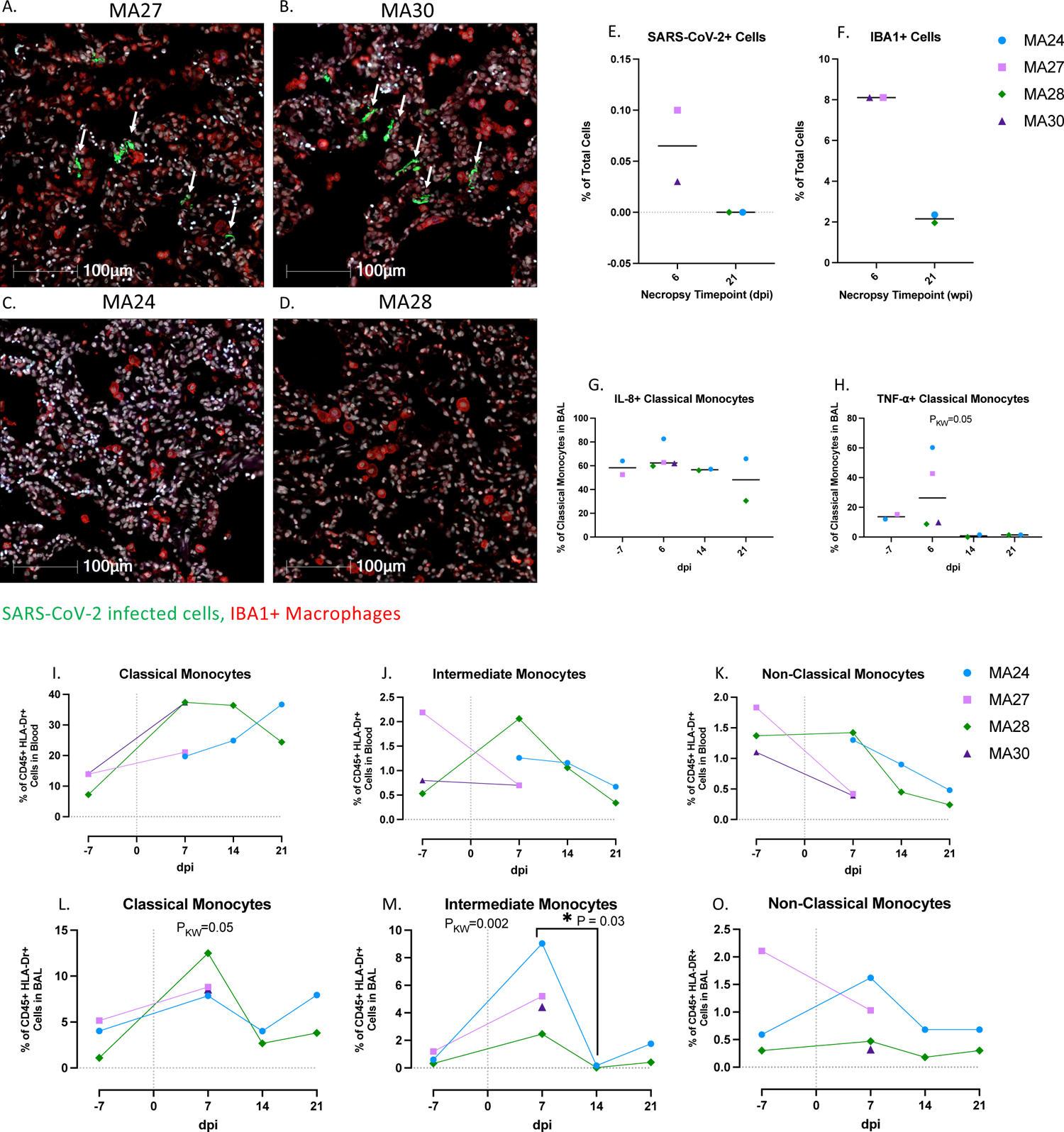
Pulmonary SARS-CoV-2 infection and macrophage/monocytes in the lung and blood. **A-D**. SARS-CoV-2 infection and macrophage infiltration in the lungs of pigtailed macaques at 6-(**A&B**) and 21-days post infection (dpi, **C&D**). DAPI=White, Green=SARS-CoV-2, Red=IBA1,Blue=Autofluorescence. **E-F.** Percentage of SARS-CoV-2 infected cells (**E**) and IBA1+ macrophages (**F**) in the lung at necropsy. Bars represent median. **G&H**. Frequency of IL-8 (**G**) and TNF-⍺ (**H**) expressing classical monocytes (CD45+ HLA-DR+ CD14-CD16+) in BAL. **I-O.** Classical (**I&L**) intermediate (CD45+ HLA-DR+ CD14+ CD16+) (**J&M**), and non-classical monocytes (CD45+ HLA-DR+ CD14-CD16+) (**K&O**) frequencies in the blood and BAL before and after SARS-CoV-2 infection. Day 0=day of infection. I-O Baseline (day-7): n=3, Day 6: n=4, Day 14: n=2, Day 21: n=2. Kruskal-Wallis comparison of overall means (P_KW_) and Dunn’s Multiple comparisons (designated by line, P_D_) tests used to determine significance. P values ≤0.05 reported

Flow cytometry showed increases in CD14+ CD16-classical monocytes in both the blood and BAL at 6-dpi (Figures 4I&L) and an increase in CD14+ CD16+ intermediate monocytes (Figure 4M) in BAL at 6-dpi. Heterogeneous fluctuations of circulating intermediate and CD14-CD16+ non-classical monocytes occurred throughout the study (Figures 4J&K). MA27 and MA24 showed increases in inflammatory cytokine, tumor necrosis factor-⍺ (TNF-⍺)-expressing classical monocytes in the BAL at 6-dpi (Figure 4H). Interleukin-1β (IL-1β) and IL-6 are key inflammatory cytokines involved in the pathophysiology of COVID-19 disease in humans^36, 37^. Here, we noted increases in IL-1β expression in peripheral CD14+ CD16-monocytes throughout the study (Figure 5A). As indicated in Figure 3A, serum levels of IL-6 remained low after infection. We also found that peripheral monocyte expression of IL-6 stayed relatively stable post infection, with only one animal, MA24, showing an increase at days 14 and 21 compared to 6-dpi (Figure 5B). There was an upward trend that was not statistically significant in neutrophil chemoattractant, IL-8, expression (Figure 5C).

**Figure 5.**
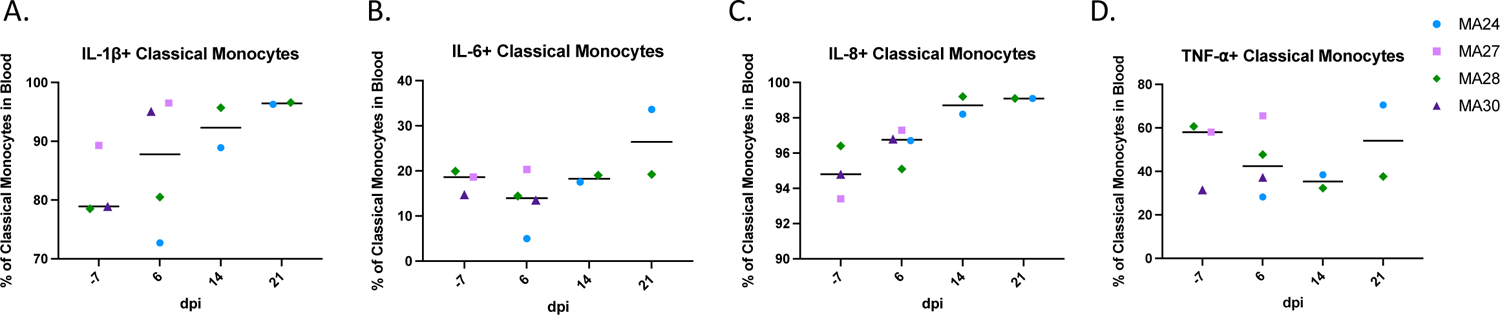
A-D. Monocyte cytokine response in the blood of pigtail macaques challenged with SARS-CoV-2. Frequency of IL-1β (**A**), IL-6 (**B**), IL-8 (**C**) and TNF-⍺ (**D**) expressing classical monocytes (CD45+ HLA-DR+ CD14-CD16+) in the blood. Bars represent median. Day 0=day of infection. Baseline (−7): n=3, Day 6: n=4, Day 14: n=2, Day 21: n=2.

### Peripheral T cell responses

Understanding the role of the adaptive immune response to SARS-CoV-2 infection is a key component to the development of effective vaccines and treatment options for COVID-19. Using flow cytometry, we measured changes to T cell populations in both the blood and BAL at baseline and 6-, 14-, and 21-dpi. CD3+ T cell fluctuations in the blood were driven by CD4 T cells which showed levels increasing significantly between days 14- and 21-pi (Figure 6A). As the percentage of CD4 T cells rise and fall over the course of the study, we observed the opposite pattern in the percentage of cytotoxic CD8 T cells (Figure 6B). We found increases in Ki-67+ CD4 T cells at 6-dpi (MA27, MA28 and MA30) and 14-dpi (MA24 and MA28) indicating increased CD4 T cell proliferation (Figure 6C&D). Increases in expression of the T cell exhaustion marker, PD-1, have been noted in a number of studies involving human COVID-19 patients^38–40^. Here we found a significant increase in PD-1+ CD4 T cells at 14-dpi (Figure 6I). Interestingly, we saw a decrease in the percentage of CD4 T cells at this same timepoint.

**Figure 6.**
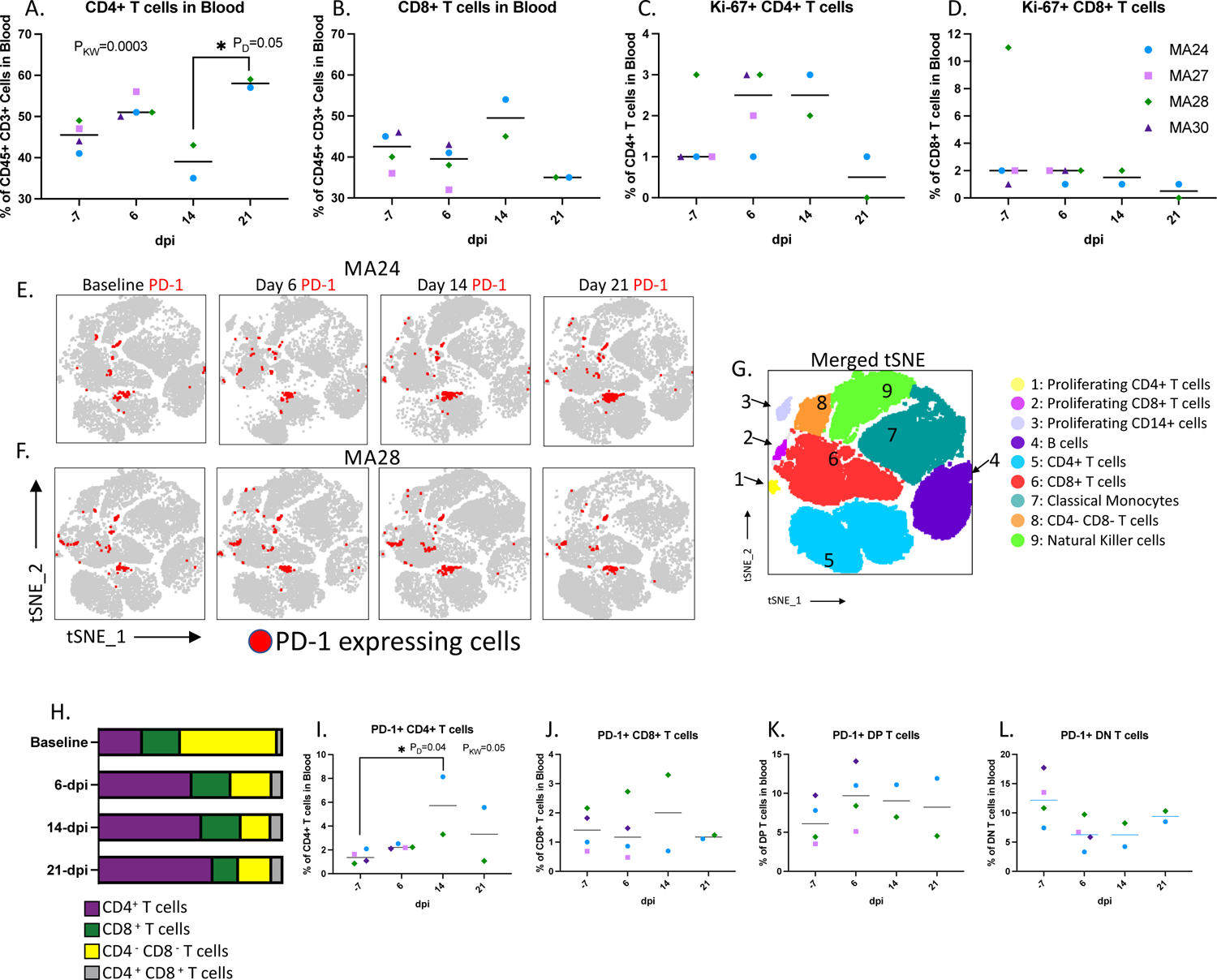
T cells in the blood. **A-B.** CD4+ (**A**) and CD8+ (**B**) T cell frequencies in the blood before and 1-, 2-, and 3-weeks post SARS-CoV-2 infection. **C-D.** Changes in Ki-67 expressing CD4+ (**C**) and CD8+ (**D**) T cells. Bars represent median **E.** tSNE plots displaying changes in PD-1 expression (red) in peripheral CD45+ cells overtime. MA24 (E) and MA28 (**F**) displayed as a representative animal. **G.** Merged tSNE indicating phenotype of the tSNE defined cell populations in E. **H.** Average changes in the percentages of CD4+, CD8+, CD4-CD8-(DN) and CD4+ CD8+ (DP) T cells within the total PD-1+ CD3+ cell population. **I-L.** Frequency of PD-1+ expressing CD4+ (**I**), CD8+ (**J**), DP (**K**) and DN (**L**) T cells in the blood. Bars represent median. Kruskal-Wallis comparison of overall means (P_KW_) and Dunn’s Multiple comparisons (designated by line, P_D_) tests used to determine significance. P values ≤0.05 reported. Baseline (−7): n=4, Day 6: n=4, Day 14: n=2, Day 21: n=2.

We next used tSNE analysis to show changes in PD-1 expressing cell populations over the course of the study (Figures 6E,F,H). At baseline, CD4-CD8-(double negative) T cells made up the greatest proportion of PD-1+ CD3+ T cells (Figure 6H). Beginning at day 6-pi, CD4 T cells made up the majority of PD-1 expressing cells, with only one animal, MA28, showing increases in PD-1 expressing cytotoxic T cells at 6 and 14-dpi (Figure 6J).

### Pulmonary T cell responses

We next sought to characterize the dynamics of pulmonary T cell populations over the course of infection by examining the frequency as well as cytokine and surface protein expression before SARS-CoV-2 infection, and at days 6-, 14-, and 21-post viral challenge. Using PMA stimulation, we noted increased frequencies of CD4+/CD8+ double positive (DP) T cells after viral challenge which remained elevated throughout the study (Figure 7A&B) ((Median DP T cells as a percentage of CD3+ T cells: Baseline: 2% (n=2), 6-dpi: 23% (n=4), 14-dpi: 30% (n=2), 21-dpi: 31% (n=2)). We also examined fold changes in surface protein and cytokine expression among the DP, CD4 and CD8 single positive T cell populations as compared to baseline in two of the animals, MA24 and MA27 (Figure 7E-J). Both MA24 (euthanized at 21-dpi) and MA27 (euthanized at 6-dpi) showed large increases among all three T cell populations in TNF-⍺ expression at 6-dpi. At fourteen days post infection, MA24 showed increased expression of Granzyme B in both the DP and CD8 T cell populations. Interestingly, it was the DP T cell population which showed the greatest fold increase in Granzyme B over baseline indicating the cytotoxic potential of this DP T cell population.

**Figure 7.**
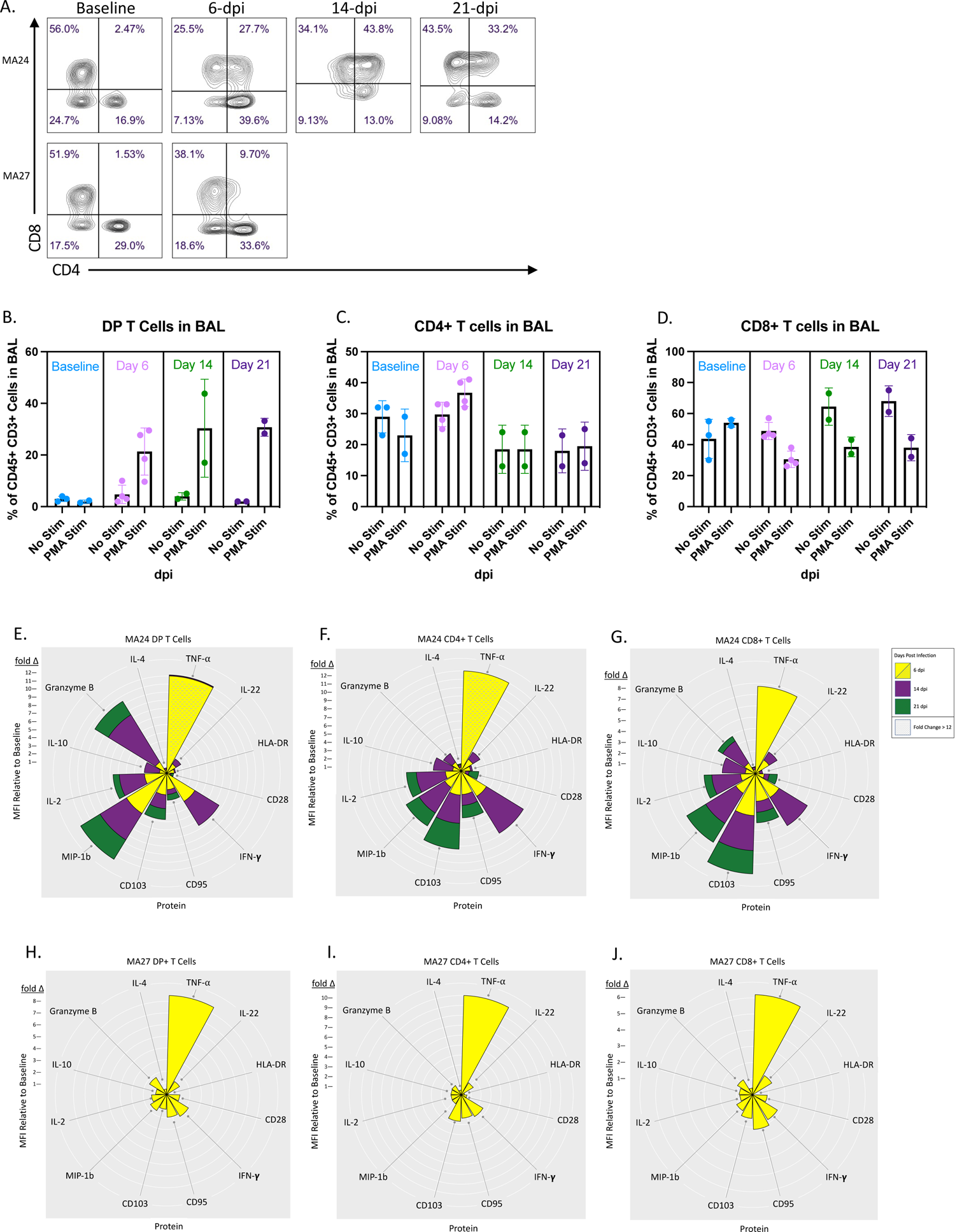
Adaptive T cell responses in the BAL. **A.** Representative flow cytometry plots showing changes in CD4 and CD8 expression in PMA/ionomycin stimulated CD3+ cells in BAL. Two animals shown (MA24 necropsied at 21-dpi, MA27 necropsied at 6-dpi). **B-D.** Effect of PMA/ionomycin on the frequency of CD4+ CD8+ (DP, **B**) CD4+ (**C**) and CD8+ (**D**) T cells in BAL before and 6-, 14-, and 21-days post SARS-CoV-2 infection. Bars represent mean and standard deviation. **E-J.** Nightingale Rose Plots (NRPs) showing fold changes in cytokine and surface protein expression compared to baseline (MFI). Yellow=6-dpi, Purple=14-dpi, Green=21-dpi. Size of petals represents magnitude of increase in expression. Distance from one white ring to the next is a 1-fold change. A decrease in expression is represented by a petal size less than the distance between two rings. Two animals shown (MA24 necropsied at 21-dpi, MA27 necropsied at 6-dpi). At 6-dpi, MA24 DP T cell TNF-⍺ MFI is 24x baseline and CD4+ T cell TNF-⍺ MFI is 50x baseline. Graph cutoff is set to a 12-fold change. **B-D.** Baseline: n=3 (No Stimulation (Stim)) and n=2 (Stim), Day 6: n=4 (No Stim) and n=4 (Stim), Day 14: n=2 (No Stim) and n=2 (Stim), Day 21: n=2 (No Stim) and n=2 (Stim).

We also compared the activity of each T cell subtype within the same time point of infection (Figure 8A-O). Prior to infection, DP T cells showed higher TNF-⍺, IFN-γ, IL-10, MIP-1β, and IL-22 expression than traditional CD4 and CD8 T cells, indicating that these cells may potentially perform a non-specific function in the pulmonary immune response^41^. After viral challenge, we found higher frequencies of Granzyme B expressing DP T cells compared to CD4 T cells and, most notably, CD8 T cells at each timepoint post infection (Figure 8E,J&O). We found significant increases in CD4 T cells expressing IL-2, IL-10 and MIP-1b (Figure 8A-E). At 14-dpi, we noted a significant increase in MIP-1β expressing CD8 T cells (Figure 8I). DP T cells also showed increased activity post viral challenge with significant increases in the frequency of IL-10, TNF-⍺, MIP-1β and Granzyme B expressing cells. Taken together, these findings show that the DP T cell population has functions which overlap with both CD4 and CD8 T cells^41^. We speculate that these cells are major histocompatibility complex class II (MHC-II) restricted CD4 T cells which upregulate CD8 upon activation, generating the described DP T cell population which has greater cytotoxic potential than traditional CD4 T cells. Pulmonary infiltrating cytotoxic CD4 T cells potentially aid CD8 T cells in viral clearance and are a unique aspect of COVID-19 disease^21^.

**Figure 8.**
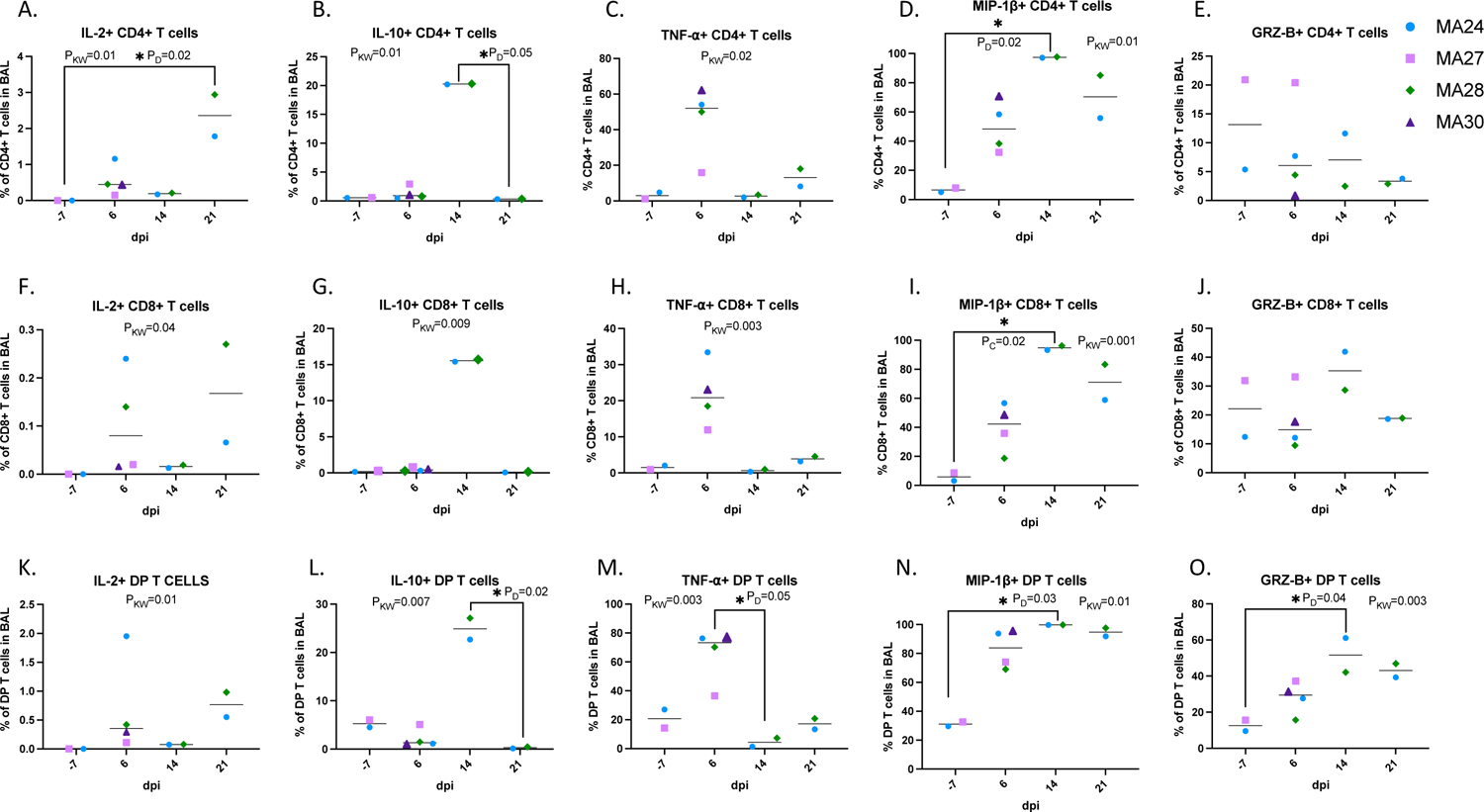
Changes in T cell cytokine expression in the lung. **A-O.** PMA/ionomycin stimulated CD4+ T cells (**A-E**), CD8+ T cells (**F-J**), and CD4+ CD8+ (DP) T cells (**K-O**). Bars represent median. Kruskal-Wallis comparison of overall means (P_KW_) and Dunn’s Multiple comparisons (designated by line, P_D_) tests used to determine significance. P values ≤0.05 reported. Baseline (−7): n=2, Day 6: n=4, Day 14: n=2, Day 21: n=2.

### CD4 T cell and Granzyme B expression in the lungs

Fluorescent Immunohistochemistry (IHC) identified cytotoxic CD4 T cells (CD4+ Granzyme B+) in the lungs of all four PTM at necropsy (Figure 9). We detected large numbers of infiltrating Granzyme B positive cells in the lungs of MA27 and MA30 (euthanized at 6-dpi) along with rare cytotoxic T cells (Figure 9A,B,F,G). At 21-dpi, MA24 (Figure 9C) showed low numbers of Granzyme B+ cells compared to MA28 (Figure 9D) and the other two animals which were euthanized at 6-dpi. Cytotoxic CD4 T cells were detected in the lung of MA28 and, with less frequency, in MA24 (Figure 9G).

**Figure 9.**
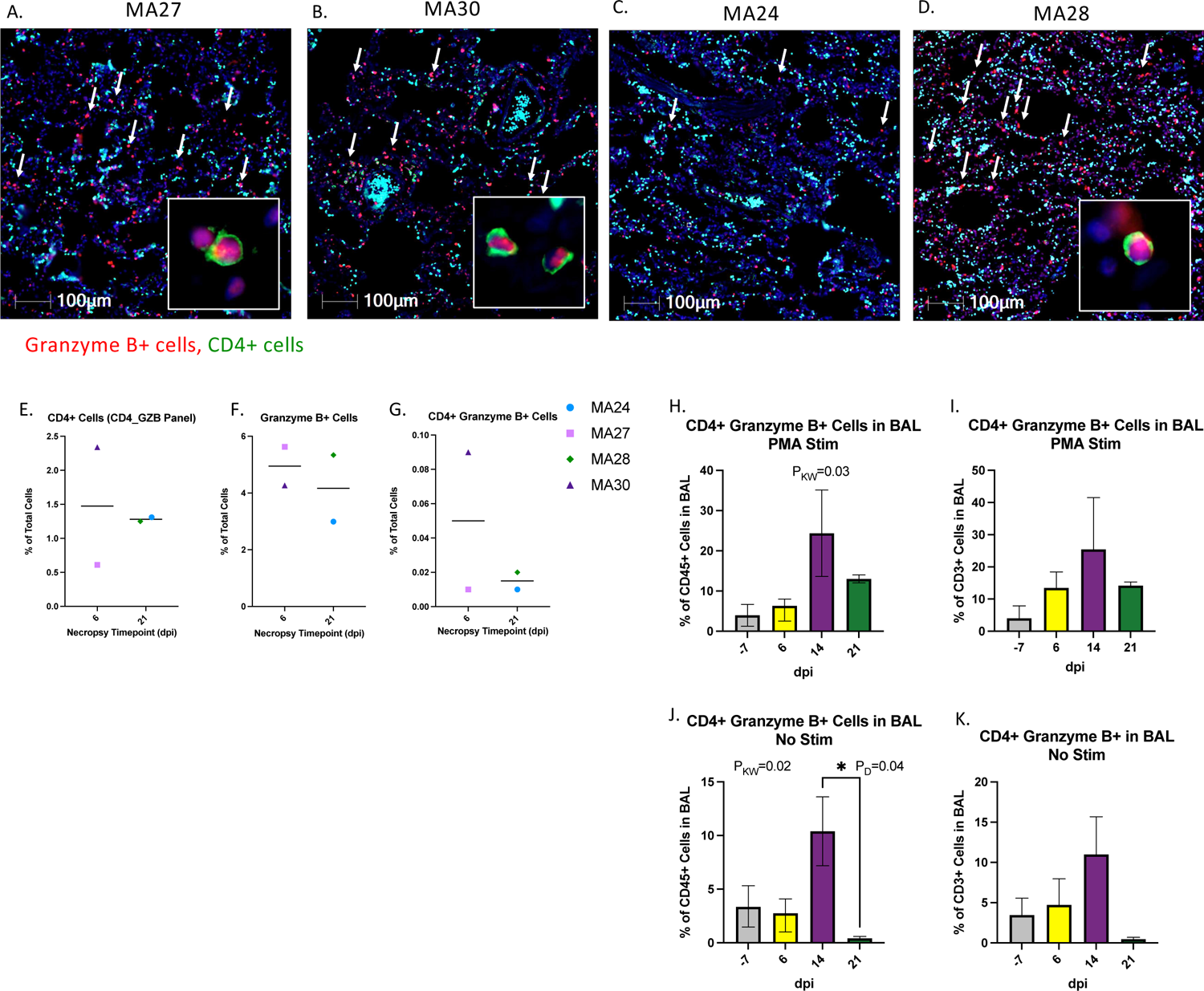
CD4 and Granzyme B expression in the lungs of SARS-CoV-2 infected macaques at 6-(A&B) and 21-days post infection (dpi, C&D). At 6-dpi, MA27 (**A**) and MA30 (**B**) the lungs are infiltrated by large numbers of Granzyme B positive cells (red, arrows). Insets: Rare CD4+ cells (green) exhibit granzyme expression. At 21-dpi, MA24 exhibits low numbers of Granzyme B positive cells (red, arrows) compared to MA28 (C) and the two, 6-dpi animals (**A&B**). DAPI=Blue, Green=CD4, Red=Granzyme B. **E-G.** Percentage of CD4+ (**E**), Granzyme B+ (**F**) and CD4+ Granzyme B+ (**G**) cells in the lung at necropsy. Bars represent median. **H-K.** CD4+ Granzyme B+ T cells (CD45+ CD3+ CD4+ Granzyme B+) in BAL at Baseline (−7) and 6-, 14- and 21-dpi as a percentage of CD45+ cells (**H&J**) and CD3+ cells (**I&K**). **H&I**. Mononuclear cells, isolated from BAL, were stimulated with PMA/ionomycin for 4-6 hours. Bars represent median and standard deviation. Kruskal-Wallis comparison of overall means (P_KW_) and Dunn’s Multiple comparisons (designated by line, P_D_) tests used to determine significance. P values ≤0.05 reported. **H-K**. Baseline: n=3 (No Stimulation (Stim)) and n=2 (Stim), Day 6: n=4 (No Stim) and n=4 (Stim), Day 14: n=2 (No Stim) and n=2 (Stim), Day 21: n=2 (No Stim) and n=2 (Stim).

We then used flow cytometry to measure cytotoxic CD4 T cells in BAL (CD45+CD3+CD4+Granzyme B+). To mirror the IHC analysis, we did not exclude CD8+ cells from our cytotoxic CD4+ population. Mononuclear cells were stimulated with PMA cocktail and cytotoxic CD4 T cells were measured as a percentage of CD45+ cells (Figure 9H&J) and CD3+ T cells (Figure 5I&K) in both the stimulated and unstimulated conditions. We noted a considerable increase in cytotoxic CD4 T cells in BAL at 14-dpi.

### SARS-CoV-2 peptide specific T cell response in the lung 21-days post infection

Mononuclear cells, isolated from BAL, were incubated overnight with SARS-CoV-2 peptides and analyzed by flow cytometry. We detected specific CD4 T cell responses against SARS-CoV-2 that localized to the lung 21 days after viral infection. Specifically, we identified CD4 T cell responses to membrane, nucleocapsid and to a lesser degree, spike peptides (Figure 10A). CD8 T cell responses against the virus were also noted, but at lower frequencies (Figure 10B). In FlowJo, we gated on the CD4 T cell population and applied tSNE analysis to identify and characterize virus specific CD4 T cells responding to membrane and nucleocapsid viral peptides (Figure 10C&D). tSNE analysis revealed a unique cluster of CD4 T cells that responded to stimulation. In this population of responding cells (Figure 10E&F), we noted increased expression of CD8 and HLA-DR, indicating cell activation. Increased expression of inflammatory cytokines and chemokines was also detected in the antiviral CD4 T cells. We noted a decrease in Granzyme B expression suggesting that the antigen specific CD4 T cells have reduced cytotoxic capacity, unlike the DP T cells (cytotoxic CD4) described previously (Figures 7, 8 and 9). As expected, antiviral CD4s have increased CD95 expression indicating a memory phenotype. Numerous studies of SARS-CoV-2 convalescent humans have described antiviral T cells with a relative predominance of CD4 T cells^42, 43^. These antiviral responses are most often noted in the blood. In our study, we were unable to detect antigen-specific T cell responses in the blood 21 days after viral infection (data not shown). Taken together, our data provide a valuable addition to the data from humans and may suggest important roles for antiviral CD4 T cells in the pulmonary compartment.

**Figure 10.**
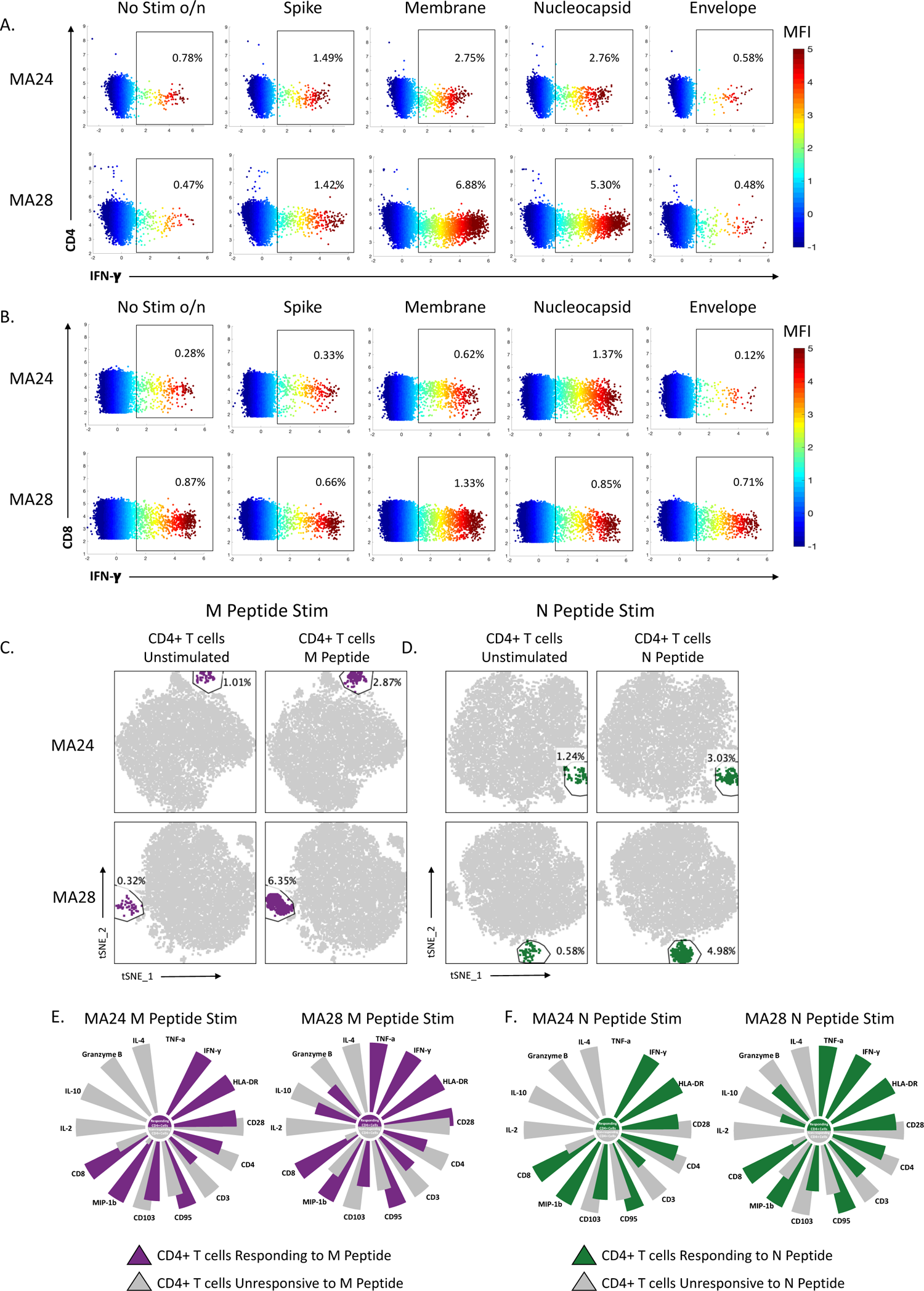
SARS-CoV-2 peptide specific T cell response in the lung 21-days post infection. Two animals shown (MA24 & MA28 necropsied 21-dpi) **A&B.** Flow cytometry dot plots showing CD4+ (**A**) and CD8+ (**B**) T cell Interferon-γ (IFN-γ) response to overnight SARS-CoV-2 peptide (spike, membrane, nucleocapsid and envelope) stimulation. No stim o/n=cells incubated overnight without stimulation. Heatmap represents arcsin transformed MFI values. **C&D.** tSNE plots of CD4+ T cells showing an expansion in cells following overnight peptide stimulation. M=SARS-CoV-2 membrane peptides (**C**), N=SARS-CoV-2 nucleocapsid peptides (**D**). **E&F.** Radial bar plot comparing MFI values of the expanded CD4+ T cell population gated on in **C&D** to the unchanged CD4+ population within the same tSNE plot. Representative animals MA24, MA28 (necropsied at 21-dpi). The higher MFI value is set to 100 and the percent difference is calculated between the higher and lower MFI values. Size of the petals represent this analysis.

### Humoral immune responses

Using flow cytometry, we measured B cell kinetics in the blood at baseline and days 6-, 14- and 21-post infection (Figure 11A). We did not detect any significant changes in the percentage of peripheral B cells over the course of the study. We next tested serum from infected animals for neutralizing antibodies using a pseudovirus assay. Not surprisingly, no neutralization was detected at 6-dpi in any sample, including the animals euthanized at that time point. By 14-dpi, neutralizing antibody responses were detectable in both MA24 and MA28 with responses decreasing by 21-dpi (Figure 11B).

**Figure 11.**
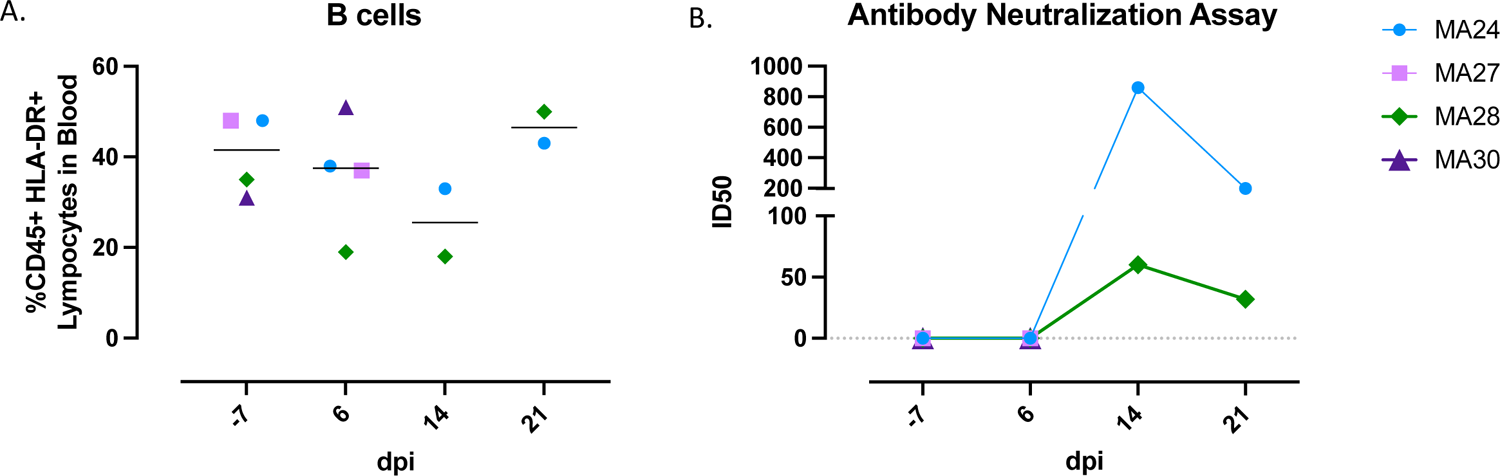
Humoral immune response in SARS-CoV-2 infected pigtail macaques. **A.** B cell frequencies in the blood before and 6-, 14-, and 21-days post (dpi) SARS-CoV-2 infection. Bars indicate median. **B.** Pseudovirus neutralization assay showing serum antibody levels against SARS-CoV-2 using HEK 293T/ACE2 cells. Baseline (−7): n=4, Day 6: n=4, Day 14: n=2, Day 21: n=2

## Discussion

The novel coronavirus SARS-CoV-2 has caused a global pandemic with little precedent. As of the time of submission, this virus has infected nearly 190 million individuals worldwide and killed four million, including over 600,000 in the United States. Illness caused by this virus, termed COVID-19, ranges from asymptomatic^44, 45^ to flu-like symptoms to severe pneumonia^46, 47^. In the most severe cases, patients have experienced acute respiratory distress syndrome (ARDS) and death^48^. It has also become apparent that a number of surprising symptoms can be associated with SARS-CoV-2 infection, including: coagulopathy, thrombosis, kidney failure and chronic respiratory/neurological issues that seemingly persist well beyond viral clearance^36, 49–56^. Although several highly effective vaccines have been created to combat the COVID-19 pandemic^57–59^, billions of individuals remain unvaccinated worldwide. Furthermore, the emergence of new viral variants with enhanced transmissibility^60–63^ and the ability to infect even the vaccinated^64^ (though this population is overwhelmingly protected from severe disease^65–67^) suggest that this virus will persist indefinitely. Barring the development and mass deployment of vaccines capable of inducing sterilizing immunity, an exceedingly difficult task, intense research focus must remain to decipher disease mechanisms so those that do become infected can be treated.

Critical to both understanding and treating the broad spectrum of disease sequelae caused by SARS-CoV-2 is the development of animal models that faithfully recapitulate COVID-19. Animal models allow timed infection and euthanasia along with extensive sample collection that are not possible during human infections. Rhesus macaques (RhM), cynomolgus macaques (CyM), and African green monkeys (AGM) have all been used to achieve this goal^12–15, 68^. To date, none of these models consistently recapitulate severe COVID-19 disease but some data suggest AGM may exhibit more severe disease than the others^12, 13^. When infected with simian immunodeficiency virus (SIV), pigtail macaques (PTM) exhibit rapid and severe disease relative to RhM and CyM, including rapid destruction of the CD4 immune compartment, severe gastrointestinal disease, and complications related to coagulopathy^17–19, 69–71^. Many of these disease features are also relevant to severe COVID-19 disease^49–54, 72–74^. A recent report demonstrated that a related species of pigtail macaques showed an abbreviated period of SARS-CoV-2 viral replication but possibly more severe disease than RhM^75^. Thus, PTM may be a reasonable model for severe disease and used to test novel therapeutics and vaccines to prevent disease.

We infected a small cohort of PTM with SARS-CoV-2 through a combination of intratracheal and intranasal instillation. Animals were tracked for viral replication in multiple sites, for immune dynamics in blood and bronchoalveolar lavage cells, and for innate and other markers of disease in blood and tissues. We identified a range of disease severity, even in our small cohort, with one animal euthanized at six days post infection showing more severe pulmonary lesions than the rest. Interestingly, multiple early indicators that are consistent with a more severe disease course in humans, were also detected in this animal, including: viral titer, an elevated neutrophil to lymphocyte ratio, elevated kynurenine to tryptophan ratio, and elevated serum inflammatory cytokines. Our findings suggest that these factors correlate with and may predict disease severity. Expanded cohort sizes that include both male and females as well as aged animals may uncover additional clinical manifestations.

Viral dynamics were similar in PTM as we have reported in RhM^12, 13^. Viral RNA, including subgenomic RNA, was consistently detected throughout the first several days of infection. We detected persistent viral titers at multiple sites in some of the animals throughout the course of the study. These data confirm PTM as a robust model of viral infection and replication, similar to RhM, and suggest this model may be used to study novel virus host relationships.

COVID-19 disease is commonly characterized by pulmonary infiltration of inflammatory immune cells^34^. Innate cells, particularly monocytes, are considered important mediators of disease progression^35^. Although infiltrating monocytes were identified in our PTM, T cells were a more dominant cellular infiltrate into lungs as detected in bronchoalveolar lavage sampling. Specifically, we identified a unique population of CD4+/CD8+ double positive T cells that upregulated inflammatory cytokines such as TNF-⍺ as well as Granzyme B over the course of infection. Traditionally, these cells would be predicted to be major histocompatibility complex class II (MHC-II) restricted CD4 T cells that upregulate CD8 upon activation. Pulmonary infiltrating CD4 T cells with cytotoxic capacity, as measured by Granzyme B, are a unique and possibly understudied aspect of COVID-19 disease.

We also identified relatively high magnitude CD4 T cell responses against the virus that localized to the lung 21 days after viral infection. CD8 T cells against the virus were also noted, but at lower frequencies. Many studies have reported antiviral T cells in SARS-CoV-2 convalescent humans, with a relative predominance of CD4 T cells, however these responses are nearly always noted in blood^42, 43^. Thus, our data provide a valuable addition to the data from humans and may suggest important roles for antiviral CD4 T cells in pulmonary sites.

Taken together, our data define a new animal model for COVID-19. PTM show robust viral replication, SARS-CoV-2 associated pneumonia, and complex innate and adaptive immune responses that may shed light on mechanisms of COVID-19 disease. This model may prove valuable for testing novel immunomodulatory therapeutics and vaccines, including those that modulate pulmonary infiltration of T cells and other inflammatory cells. Finally, our data confirmed COVID-19 associated inflammation was not always resolved 21dpi, despite no evidence of continued viral replication at that time point. Thus, this model may also be valuable for the study of long-term chronic effects associated with SARS-CoV-2 infection.

## Acknowledgements

The following reagents were obtained through BEI Resources, NIAID, NIH: Peptide arrays NR4219, NR-52402, NR-53822, NR-53822. Funding was provided by NIH NAID grants P51OD01110459, R21 AI150413-01, R01 AI38782-01, and R24 AI120942.

---

**Supplemental Figure 1.**
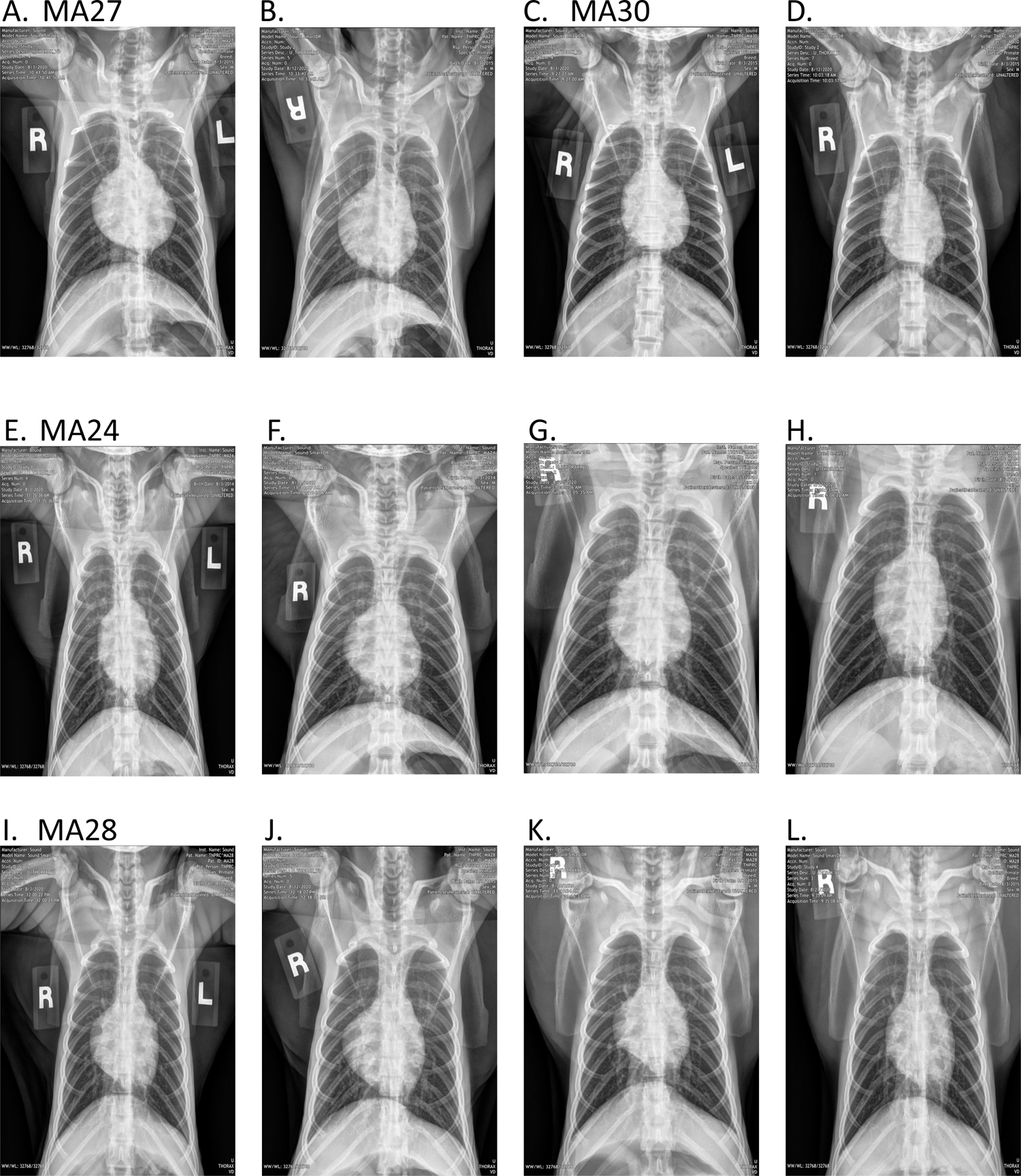
Radiographs of pigtail macaques (PTM) challenged with SARS-CoV-2. MA27 baseline (**A**) and 6-days post infection (dpi) (**B**). MA30 at baseline (**C**) and 6-dpi (**D**). MA24 at baseline (**E**), 6-dpi (**F**), 14-dpi (**G**) and 21-dpi (**H**). MA28 at baseline (**I**), 6-dpi (**J**), 14-dpi (**K**) and 21-dpi (**L**). Baseline for all four PTM was established 3-days prior to infection.

**Supplemental Figure 2.**
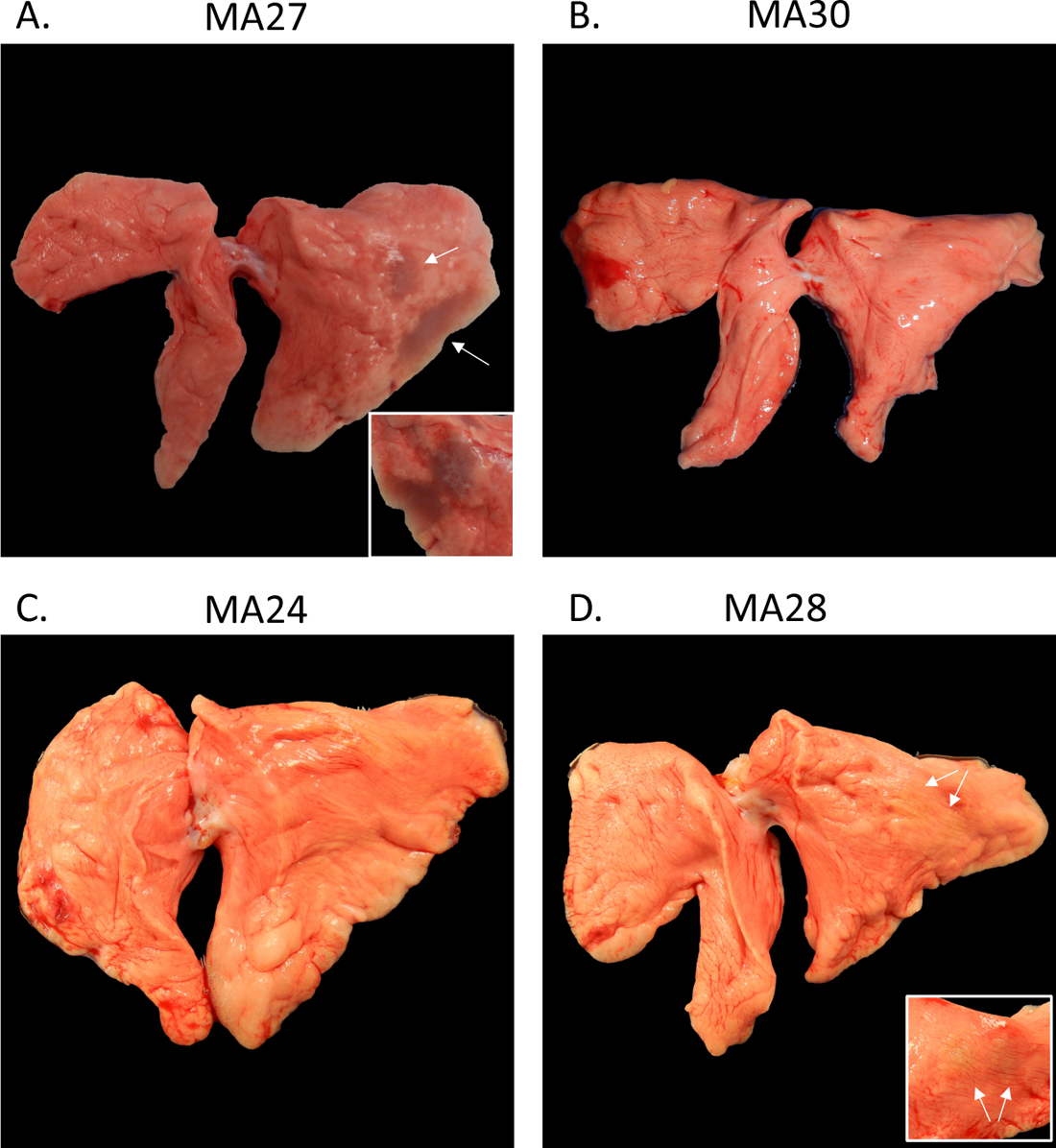
Gross pathological pulmonary pathology in SARS-CoV-2 infected pigtail macaques (PTM). **A-D.** Gross pulmonary pathology at 6-(**A&B**) and 21-days post infection (dpi, **C&D**). **A**. MA27, the left caudal lung lobe has multifocal tan-plum areas of consolidation (arrows). Inset: the consolidation extends to the diaphragmatic and medial surface of the left caudal lung. There is no evidence of gross pathology in MA30 (B) or MA24 (C). **D.** MA28, the laterodorsal aspect of the left caudal lobe contains two small, flat tan foci (arrows). Inset: closer view of tan foci.

**Supplemental Figure 3.**
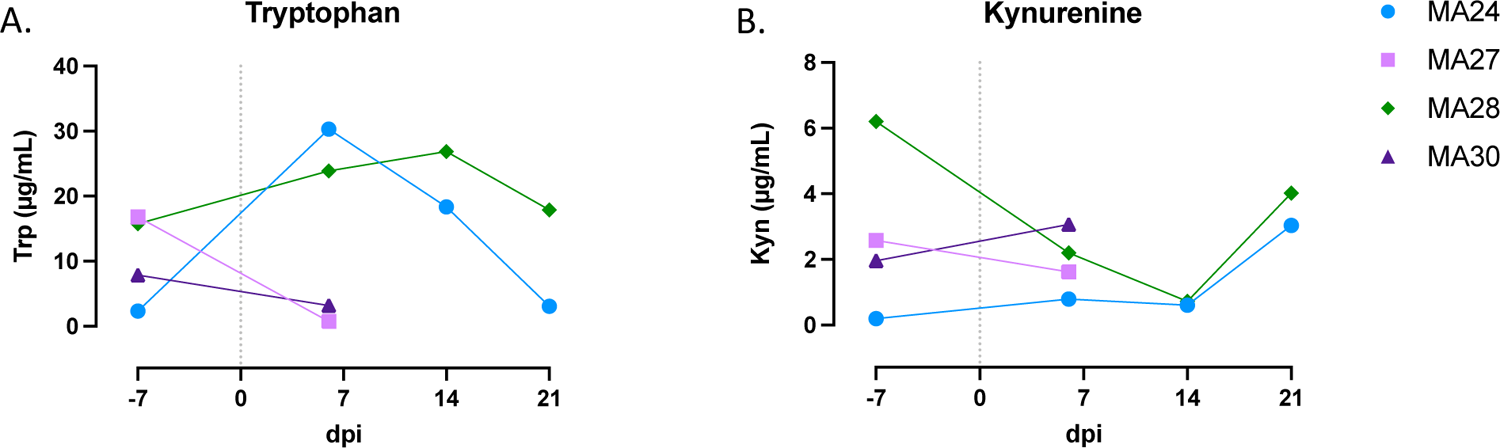
Changes in IDO activity post SARS-CoV-2 Infection. **A.** Tryptophan (Trp) and **B.** Kynurenine (Kyn) levels in plasma before and after SARS-CoV-2 infection. Day 0=day of infection, Baseline: n=4, Day 7: n=4, Day 14: n=2, Day 21: n=2. dpi=days post infection.

**Supplemental Table 1.**
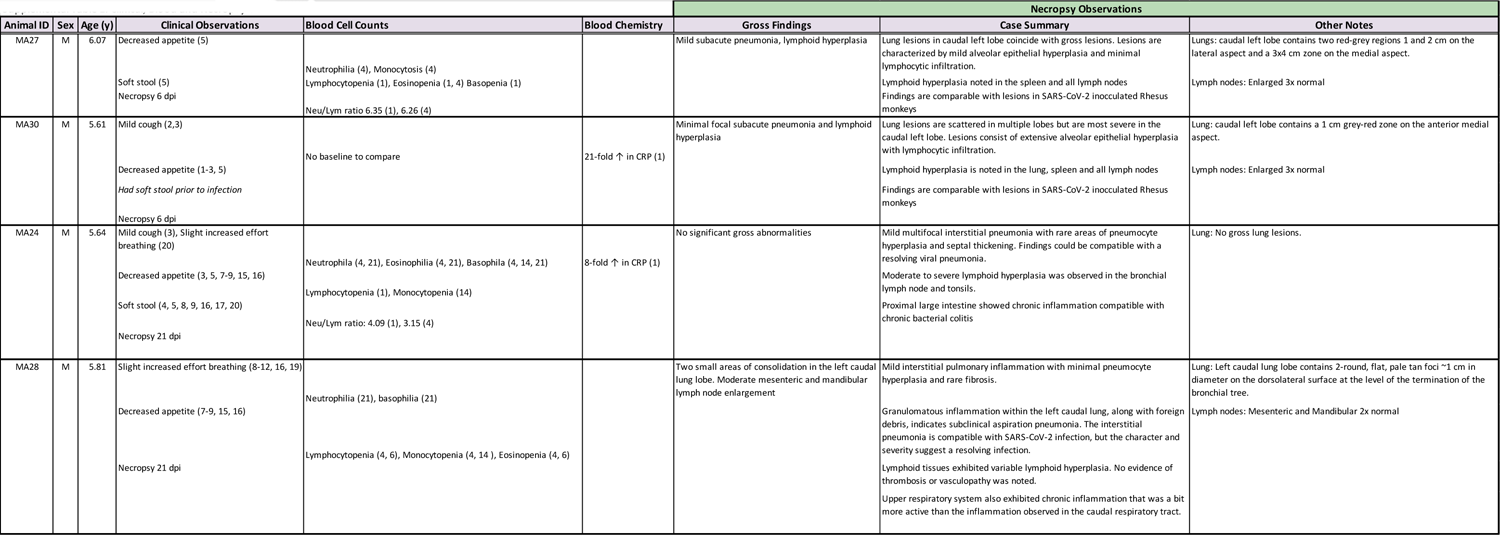
Clinical, blood and necropsy observations in pigtail macaques (PTM) challenged with SARS-CoV-2 (n=4). Values in parenthesis represent days post SARS-CoV-2 Infection. Neutrophilia, eosinophilia, basophilia, monocytosis defined as ≥ 2-fold increase over baseline^16^. Lymphocytopenia, monocytopenia, eosinopenia, basopenia defined as a 35% reduction from baseline^16^. CRP: C-reactive protein

**Supplemental Table 2.** Monocyte Flow Cytometry Panel.

| Supplemental Table 2. Monocyte Panel |  |  |  |  |  |  |
| --- | --- | --- | --- | --- | --- | --- |
| Fluorochrome | Antigen | Volume (uL)/Test | Clone | Catalog# | Lot# | Company |
| FITC | IL-6* | 5 | MQ2-6A3 | 554696 | 6168973 | BD Pharmingen |
| PCP-Cy5.5 | CD169 | 5 | 7-239 | 346020 | B309159 | BioLegend |
| AL647 | IL-1B* | 5 | JK1B-1 | 508208 | B274564 | BioLegend |
| AL700 | TNF-a* | 5 | MAb11 | 502928 | B221571 | BioLegend |
| APC-Cy7 | HLA-DR | 5 | L243 | 307618 | B295253 | BioLegend |
| PacBlue | CD66 | 2 | TET2 | 130-119-851 | 1320070484 | Miltenyi Biotec |
| BV510 | L/D | 0.5 | Fixable Aqua Dead Cell Stain Kit | L34957 |  | Invitrogen |
| BV605 | CD45 | 5 | D058-1283 | 564098 | 9051992 | BD Horizon |
| BV650 | CD11c | 5 | S-HCL-3 | 744437 | 0203626 | BD OptiBuild |
| BV711 | CD16 | 5 | 3G8 | 302044 | B296475 | BioLegend |
| BV786 | CD103 | 5 | Ber-ACT8 | 350230 | B274642 | BioLegend |
| PE-CF594 | IL-8* | 5 | G265-8 | 563531 | 0219387 | BD Horizon |
| PE-Cy7 | CD68* | 5 | Y1/82A | 565595 | 0079803 | BD Pharmingen |
| BUV395 | CD123 | 5 | 7G3 | 564195 | 9337379 | BD Horizon |
| BUV496 | CD206 | 5 | 19.2 | 741173 | 0261803 | BD OptiBuild |
| BUV737 | CD14 | 5 | M5E2 | 612764 | 0140605 | BD Horizon |

**Supplemental Table 3.** Phenotype Flow Cytometry Panel.

| Supplemental Table 3. Phenotype Panel |  |  |  |  |  |  |
| --- | --- | --- | --- | --- | --- | --- |
| Fluorochrome | Antigen | Volume (uL)/Test | Clone | Catalog# | Lot# | Company |
| FITC | CD21 | 5 | B-Ly4 | 561372 | 0149866 | BD Pharmingen |
| PCP-Cy5.5 | CD169 | 5 | 7-239 | 346020 | B309159 | BioLegend |
| APC | CD11c | 5 | S-HCL-3 | 340714 | 70397 | BD Pharmingen |
| AL700 | Ki67* | 5 | B56 | 561277 | 9261927 | BD Pharmingen |
| APC-Cy7 | CD20 | 5 | L27 | 561277 | 8320624 | BD Bioscience |
| PacBlue | CD8 | 5 | SK1 | 344718 | B280004 | BioLegend |
| BV510 | L/D | 0.5 | Fixable Aqua Dead Cell Stain Kit | L34957 |  | Invitrogen |
| BV605 | CD4 | 5 | OKT4 | 317438 | B297647 | BioLegend |
| BV650 | CD3 | 5 | SP34-2 | 563916 | D163859 | BD Horizon |
| BV711 | CD16 | 5 | 3G8 | 302044 | B296475 | BioLegend |
| BV785 | CD27 | 5 | O323 | 302832 | B264783 | BioLegend |
| PE | FasL | 5 | NOK-1 | 306407 | B269886 | BioLegend |
| PE-CF594 | HLA-DR | 5 | L243 | 307654 | B320176 | BioLegend |
| PE-Cy5 | CD95 | 5 | DX2 | 15-0959-42 | 2252537 | Invitrogen |
| PE-Cy7 | PD-1 (CD279) | 5 | J105 | 25-2799-42 | 228650 | Invitrogen |
| BUV395 | CD45 | 5 | D058-1283 | 564099 | 0192257 | BD Horizon |
| BUV496 | CD28 | 5 | CD28.2 | 741168 | 0203628 | BD OptiBuild |
| BUV737 | CD14 | 5 | M5E2 | 612764 | 0140605 | BD Horizon |

\* Intracellular

**Supplemental Table 4.** T cell Flow Cytometry Panel.

| Supplemental Table 4. T cell Panel |  |  |  |  |  |  |
| --- | --- | --- | --- | --- | --- | --- |
| Fluorochrome | Antigen | Volume (uL)/Test | Clone | Catalog# | Lot# | Company |
| FITC | MIP-1b* | 20 | D21-1351 | 560565 | 348912 | BD Parmingen |
| PCP-Cy5.5 | IL-2* | 5 | MQ1-17H12 | 560708 | 8142599 | BD Parmingen |
| AL647 | IFN-g* | 5 | 4S-B3 | 502516 | B228928 | BioLegend |
| AL700 | TNF-a* | 5 | MAb11 | 557996 | 8179967 | BD Parmingen |
| APC-Cy7 | HLA-DR | 5 | L243 | 307618 | B295253 | BioLegend |
| PacBlue | CD8 | 5 | SK1 | 344718 | B280004 | BioLegend |
| BV510 | Live/Dead | 0.5 | Fixable Aqua Dead Cell Stain Kit | L34957 |  | Invitrogen |
| BV605 | CD4 | 5 | OKT4 | 317438 | B297647 | BioLegend |
| BV650 | CD3 | 5 | SP34-2 | 563916 | D163859 | BD Horizon |
| BV711 | CD95 | 5 | DX2 | 563132 | 004458 | BD Horizon |
| PE | IL10* | 5 | JES3-9D7 | 501404 | B273796 | BioLegend |
| PE-CF594 | GZB* | 5 | GB11 | 562462 | 0050626 | BD Horizon |
| PE-Cy7 | IL-4* | 5 | MP4-25D2 | 500824 | B293168 | BioLegend |
| BUV395 | CD45 | 5 | D058-1283 | 564099 | 0101778 | BD Horizon |
| BUV496 | CD28 | 5 | CD28.2 | 741168 | 0203628 | BD OptiBuild |

\* Intracellular

**Supplemental Table 5.** T cell SARS-CoV-2 Peptide, PMA/Ionomycing Stimulation

| Supplemental Table 5. T cell SARS-CoV-2 Peptide, PMA/Ionomycin Stimulation |  |  |  |  |
| --- | --- | --- | --- | --- |
| Stimulant | Volume (uL)/mL | Catalog# | Lot# | Company |
| Cell Stimulation Cocktail | 1 | 423301 | B295704 | BioLegend |
| SARS-CoV-2 Spike Peptide | 10 | NR-52402 |  | BEI Resources, NIAID, NIH |
| SARS-CoV-2 Membrane Peptide | 5 | NR-53822 |  | BEI Resources, NIAID, NIH |
| SARS-CoV-2 Nucleocapsid Peptide | 5 | NR-52419 |  | BEI Resources, NIAID, NIH |
| SARS-CoV-2 Envelope Peptide | 5 | NR-53822 |  | BEI Resources, NIAID, NIH |
| Brefeldin-A | 1 | 420601 | B211242 | BioLegend |

**Supplemental Table 6.** Immunohistochemistry Reagent Panel.

| Supplemental Table 6. Immunohistochemistry |  |  |  |  |  |  |
| --- | --- | --- | --- | --- | --- | --- |
| Primary Antibody | Species | Company | Catalog# | Dilution | Secondary Antibody |  |
|  |  |  |  |  | Species | Fluorochrome |
| SARS | Guinea Pig | BEI | NR-10361 | 1:1000 | Goat anti-guinea pig | Alexa Fluor488 |
| IBA1 | Rabbit | Wako | 019-19741 | 1:50 | Goat anti-Rabbit | Alexa Fluor568 |
| CD3 | Mouse IgG1 | Agilent | M7254 | 1:20 | Goat anti-MS IgG1 | Alexa Fluor568 |
| CD4 | Rabbit | Abcam | Ab133616 | 1:10 | Goat anti-Rabbit | Alexa Fluor488 |
| Granzyme | Mouse IgG2a | Agilent | M7235 | 1:50 | Permanent Red (Agilent K0640) |  |
| Dapi |  | Invitrogen | D1306 | 1:20,000 |  |  |

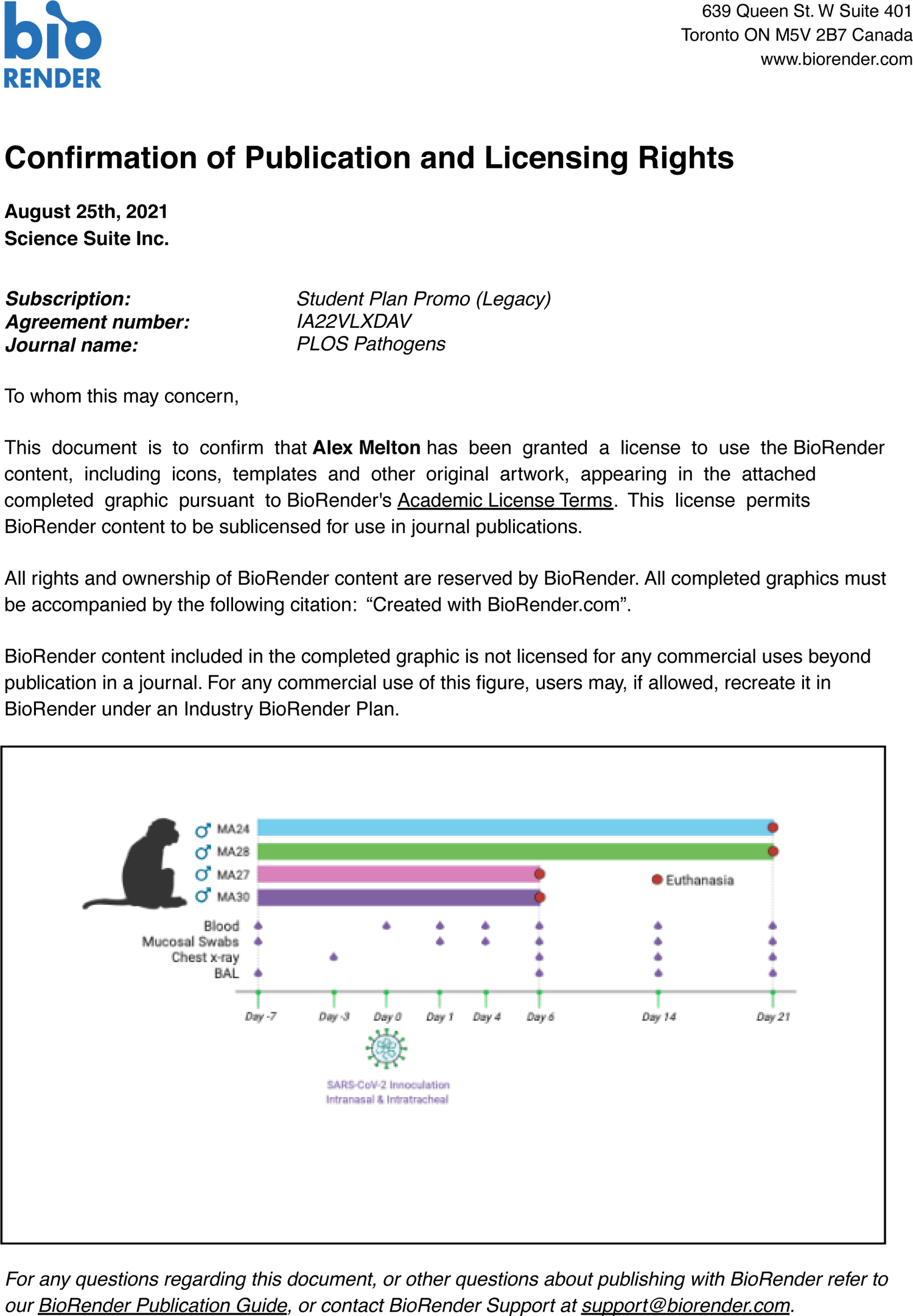

